# A model of the locust visual system under diverse stimuli highlights functionality of various mechanisms and suggests a parsimonious structure

**DOI:** 10.64898/2026.09.14.751446

**Authors:** Erik GN Olson, Travis K Wiens, John R Gray

## Abstract

Locusts possess a collision-sensitive pathway in their visual system culminating in a neuron, the lobula giant movement detector (LGMD), which preferentially responds to looming stimuli. The LGMD is also notable for its reduced or absent response to other stimuli, such as translating objects, wide-field motion, and incoherent images of looming objects. While experimental and modelling work have both shed light on the various mechanisms underpinning these properties, broad examinations of how these mechanisms interact with each other and with diverse stimulus types have been limited. To address this, we developed a model incorporating mechanisms from recent literature, and subjected to an array of looming stimuli with varying size-to-speed ratio, polarity (OFF and ON), and coherence; wide-field background motion, translating stimuli, and trajectory changes were also examined. The model showed quantitative and qualitative fidelity to biological data in its replication of a wide variety of stimulus responses, with its preference for incoherent visual stimuli being a notable exception. Based on investigations with various inhibition types removed, it was shown that lateral inhibition predominantly suppressed wide-field motion responses. Global inhibition both normalized growing excitation over the course of looming, and improved recognition of looming stimuli against moving backgrounds. Feedforward inhibition was found to have diverse roles, including shaping peak response time and its variability, and improving coherence selectivity. The success of the model in replicating multiple stimuli also showed the plausibility of several underlying assumptions, including sharing of input between the LGMD and multiple other neuron types.

**Author summary:** The locust LGMD is a key visual neuron that detects looming objects approaching on a collision course. We modelled this neuron and the network of neurons associated with it to test hypotheses about the structure of the network, and to better understand functions of its various parts. Our hypotheses included that the neurons which excite the LGMD and those which inhibit it share the same inputs, keeping the network simpler. Both the LGMD and other neurons in our model behaved similarly to real life in multiple situations, supporting our hypothesis. By examining the model LGMD outputs with various mechanisms removed from the model, we determined the roles and functions of these mechanisms in recognizing and responding to impending collisions. Two separate mechanisms—global and lateral inhibition—were required to detect collisions when a moving background was present, a task vital to a locust in flight. Another mechanism—feedforward inhibition—showed potential to make escapes from predators more timely but less predictable, and to help in distinguishing true colliding objects from similar background noise. Overall, our model furthers understanding of how locusts visually detect and respond to collisions, and provides a useful platform for future work.

## Introduction

The dual tasks of locating oncoming objects and avoiding them are key functions of animal visual and motor systems, respectively. A particularly well-studied example of a neuron capable of responding to approaching obstacles and effecting responses to them is that of the locust descending contralateral movement detector (DCMD). These neurons make up a bilaterally symmetric pair, and carry signals encoding the imminence of collision [1–3] to the motor nervous system of the animal [4]. Firing of the DCMD neurons has been correlated with escape jumps [5], evasive gliding [6–8] and steering [9, 10] behaviours in flying locusts.

While these neurons provide visual data to motor centers, their role is primarily that of transmission; the activity of the DCMD neurons follows that of their presynaptic partner, the lobula giant movement detector (LGMD), with a one-to-one correspondence of spikes [11]. The LGMD, or more precisely the LGMD1, is one of a pair of neurons within the lobula region of the locust’s optic lobe which responds preferentially to looming objects [12] approaching the eye at a constant velocity. The LGMD1 exhibits sensitivity to visual inputs containing light-to-dark (i.e. OFF) and dark-to-light (i.e. ON) stimuli, while the LGMD2 exhibits sensitivity only to OFF stimuli [12]. Given that only the LGMD1 is known to make connections with the DCMD, it is generally the more frequently studied of the two LGMDs. This is likely due to both the relative ease of recording from the DCMD neuron, and the greater implications for locust behaviour. As such, the term “LGMD” generally refers to the LGMD1, and will be used as such here.

The LGMD is a large, electrotonically extensive neuron [13], with different portions of its structure performing different computational functions. It possesses three large dendritic subfields, each of which receives different inputs [14]:

- field A, which receives OFF and ON excitation from the trans-medullary afferents (TmAs) via the second optic chiasm;
- field B, which receives feedforward inhibition relating to ON stimuli via the second optic chiasm; and
- field C, which receives input via the dorsal uncrossed bundle (DUB).

Field C was classically understood to also receive feedforward inhibition, albeit from OFF stimuli [14, 15]; however its role has recently been expanded [16], as discussed below. This structural complexity of the LGMD is paired with a great number of neurophysiological mechanisms—both within the LGMD and within its presynaptic network—which shape not only the LGMD’s responses to looming stimuli, but its responses (or lack thereof) to numerous other stimuli as well.

### Stimulus responses of the LGMD, and underlying physiology

Hatsopoulos et al. [17] first recognized that the looming responses of the LGMD generally follow the *η* function, which Jones and Gabbiani [18] presented in the form

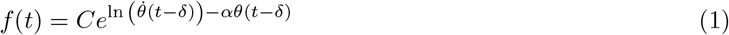

where *f* (*t*) is the firing rate of the LGMD [spikes/s] at time *t* [s]. *θ*(*t* − *δ*) is visual subtense angle of the looming object [rad] subjected to a fixed delay *δ* [s], and 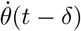 is the object expansion rate [rad/s] subjected to the same delay. *C* and *α* are constants governing the shape of the firing rate curve. This function makes several predictions that are true of the LGMD and the network of neurons providing input to it. These include

- that excitatory input to the LGMD is tied to the angular velocity of looming object edges [19],
- that excitation is transformed in a sublinear, approximately logarithmic fashion [18];
- that inhibitory input to the LGMD follows the angular size of the looming object [15];
- that the transformation of summed excitation and inhibition into firing rate constitutes an approximately exponential relationship [20];
- that firing of the LGMD peaks prior to collision [1, 6, 17]; and
- that the latency between this peak time and collision increases linearly with the size-to-speed ratio of looming, *l/*|*v*| [1, 17, 21].

As the latter two points depend on the physiological elements of the prior ones, testing an LGMD model for a linear progression of peak time versus *l/* |*v*| provides a useful point of reference for its validity. However, replicating this alone is not enough, as it is possible to develop models which exhibit such behaviour (e.g. the model of Bermúdez i Badia et al. [22]) while incorporating elements which are distinctly lacking from the LGMD and its associated neuroanatomy (e.g. elementary motion detectors; see Jones and Gabbiani [19]). As such, an effective model must have clear correspondences with anatomy, and also requires further points of reference for its behaviour.

The LGMD is not only sensitive to looming, but exhibits a preference for it over other object motions such as translation [23, 24] and recession [25]. One mechanism shaping this preference is spike frequency adaptation, enabled by calcium dependent potassium conductances [26]. These inhibitory conductances build up over extended firing, penalizing responses to stimuli which remain constant in intensity, such as translation. In contrast, the rapidly increasing excitation generated by looming allows increasing firing rates.

Regardless of mechanism, the responses of the LGMD to translation and looming are distinctly different, but the LGMD can rapidly switch to a new, appropriate response as an object’s trajectory changes [23], as well as being able to adjust its response to changes in loom approach speed [1]. However, instantaneous changes in the trajectory of object motion are often accompanied by brief transient changes in firing rate before the LGMD switches to encoding the new motion, with observations including

- brief drops in firing rate after transitions from translating to looming [23, 24, 27]; and
- brief increases in firing rate after transitions from looming to translating [23].

While decreasing response after a switch to a preferred stimulus (or vice-versa) may seem counterintuitive, it may be meaningful given the relationships observed between dropping LGMD firing rate and triggering of escape behaviour [5]. Regardless of its biological purpose, it provides another useful point of reference for computational models seeking to capture a broad spectrum of LGMD behaviour.

The LGMD is distinguished not only by what it responds to, but also what it does not. Sustained wide-field motion in the visual field, such as might be generated by the animal’s own motion—and used as a visual cue for guidance during flight [28, 29]—does not trigger a sustained response at the LGMD [27, 30]. However, the LGMD can still distinguish and respond to looming and translating stimuli when background motion is present throughout the visual field [27], albeit with delays to the onset and peak of firing. The suppression of this wide-field motion is classically attributed to lateral inhibition presynaptic to the LGMD [30]. The neurons of the DUB also show a similar lack of sustained response to wide-field motion [15]; modelling work by Olson et al. [31] showed that this may also be due to similar lateral inhibition.

Another key ability of the LGMD is to distinguish looming stimuli from those producing similar changes in luminance, but not a coherent image. The LGMD responds more strongly to stimuli showing a coherent looming object over those with equivalent luminance, but spatially incoherent mapping that does not form a contiguous object [16, 32, 33]. Two complementary mechanisms have been documented which implement this coherence selectivity, both of which depend on the fact that excitation to field A of the LGMD is retinotopically mapped [34, 35]. The first is lateral excitation between the TmAs providing excitation to field A of the LGMD. Each TmA branches near the LGMD, with the resulting terminals not only making synapses with the LGMD, but also reciprocally with other TmA terminals [36, 37]. These reciprocal synapses are excitatory, and increase the strength of synaptic inputs from the TmAs to the LGMD [33]. This arrangement selectively increases LGMD excitation when stimuli are spatially contiguous.

The second mechanism of coherence selectivity are active conductances within field A of the LGMD [32]—namely, hyperpolarization-activated cyclic nucleotide-gated nonselective cation (HCN) conductances, and potassium conductances resembling K_D_ channels, termed K_D-like_ by Dewell and Gabbiani [32]. These conductances work in tandem, and respond to localized excitation and depolarization by closing, which

- decreases local membrane conductance;
- increases local membrane time constant; and
- enhances integration of excitatory inputs.

Non-localized excitation from incoherent stimuli does not produce sufficient localized depolarizations to close these voltage-gated channels, resulting in reduced integration of incoherent synaptic input, and thus coherence selectivity from the LGMD.

The above mechanisms of coherence selectivity apply predominantly to OFF looms; that is, dark stimuli approaching against a light background. LGMD responses to ON looms (light objects against a dark background) lack the distinct coherence selectivity of their OFF counterparts [16]. Dewell et al. [16] found that this was likely the result of large portions of ON excitation arriving not at field A of the LGMD, but at field C via a subgroup of neurons in the DUB—a discovery which substantially revises classical understanding of both the DUB and dendritic field C. These field C excitatory inputs lacked the retinotopic mapping needed for the mechanisms of coherence selectivity implemented in field A, which appears to explain the lack of coherence selectivity in LGMD responses to ON looms.

Two other important mechanisms not related to any of the above stimulus types specifically are global inhibition, and M currents. Global inhibition was identified in the inputs to the LGMD by Zhu et al. [33], and appears to reduce excess firing in the latter part of a looming object’s approach, allowing an expansion of the LGMD’s dynamic range and responses to earlier portions of the loom. Olson et al. [31], via modelling, showed that similar global inhibition may shape the responses of DUB neurons which provide feedforward inhibition to the LGMD (corresponding to the *θ*(*t* − *δ*) term in Eq 1). M currents are potassium channels which [38] found to shape the firing behaviour of the LGMD. Namely, they appear to influence whether spikes are fired singly, or in high-frequency bursts—something which changes prominently over the time course of a looming stimulus [39].

While many of the physiological mechanisms underlying LGMD stimulus responses described above have been firmly identified and localized within the locust visual system, many have not—particularly those occurring prior to the LGMD and its immediate surroundings. Based on the observations of DUB neuron behaviour by Wang et al. [15] and DUB neuron dendritic inputs by Rowell et al. [14], Olson et al. [31] posited that global and lateral inhibition were located in a portion of the optic lobe forming shared inputs to both the LGMD and the DUB neurons, up to the point of the TmAs which excite field A of the LGMD forming the inputs of DUB neurons. However, they did not confirm that their modelling of these inputs would also provide appropriate stimulus to an LGMD neuron, restricting the scope of their investigation to the outputs of DUB neurons. They also did not investigate the relative localization of global and lateral inhibition within the optic lobe.

In general, the scope of LGMD models in the literature to this point has been limited, with particular models primarily considering a small subset of physiological aspects and LGMD responses at a time; recent examples include models examining

- localization of excitation between fields A and C, and looming responses [16];
- field A active conductances and coherence selectivity [32];
- M currents and bursting during looms [32];
- field A structure and looming responses [18]; and
- calcium dependent potassium conductances, and looming and translating responses [26].

While investigations such as these have been vital for deepening understanding of the physiology and stimuli in question, an investigation of an LGMD model covering a wide variety of stimuli and computational mechanisms would enable a different set of insights. In particular, such an investigation would allow

- examination of how different mechanisms in the LGMD and its inputs interact with multiple stimuli and with each other;
- confirmation that the implementations of these mechanisms reproduces a wide variety of responses, increasing the likelihood that these implementations are biologically meaningful; and
- development of an LGMD model that can simulate in vivo responses to complex visual environments incorporating diverse stimuli.

### Objectives and assumptions

The primary objectives of this work are twofold. The first is to use a dynamic computational model to examine how the diverse elements identified experimentally within the LGMD and its inputs might shape a wide variety of stimulus types, both individually and in concert with each other. The elements incorporated into the model include, but are not limited to:

- lateral inhibition;
- global inhibition;
- feedforward inhibition to the LGMD;
- active conductances in the LGMD such as HCN, K_D-like_, and M currents; and
- differing localization of ON and OFF excitation to different dendritic fields. The stimuli covered include

- OFF looms across varying *l/*|*v*|;
- ON looms;
- coherent and incoherent looms, both OFF and ON;
- looms with and without wide-field motion as a background; and
- stimuli incorporating looming, translating, and combinations of the two.

These were chosen to enable both quantitative and qualitative evaluation of model responses against experimental data.

The second objective of this work is to use the performance of the model across the diverse stimuli employed to examine the plausibility of several underlying neuroanatomical assumptions. The first is that the LGMD and the feedforward inhibitory neurons of the DUB share the same input network, up to and including the TmA neurons. This idea was initially explored by Olson et al. [31], which only showed that computational motifs may be in common between their inputs; this work will examine whether it is plausible for these neurons to directly share their inputs.

As an expansion of the ideas put forward by Olson et al. [31], we also explored whether the excitatory neurons of the DUB (as identified by Dewell et al. [16]) could plausibly receive their inputs directly from the TmAs as well. As these neurons are predominantly (but not completely) ON sensitive [16], while the feedforward inhibitory neurons in the DUB are OFF sensitive [14], such a scheme would require the ability to receive separate ON and OFF inputs from the TmAs. This leads to a third neuroanatomical possibility to be examined: that there exist separate OFF and ON TmAs.

The separation of the TmAs feeding excitation to the LGMD into separate ON and OFF channels runs contrary to classical thought—based on observations by O’Shea and Rowell [40]—that the TmAs are ON/OFF cells. The observations underpinning this separation are not all new, however; Strausfeld and Nässel [41] observed as early as 1981 that two distinct patterns of branching existed in the inputs to field A of the LGMD—suggesting two different types of neuron—and that the number of afferents to field A totalled 15,000—which would be consistent with two separate afferent paths per ommatidium given the roughly 7,500 to 8,000 ommatidia present in the locust eye [42, 43]. Multiple additional factors strengthen the idea that there are two separate TmA populations which, if not entirely segregated, are at least more strongly sensitive to ON and OFF signals, respectively. Recent observations in favour of this include

- the recent discovery by Wernitznig et al. [37] of two distinctive types of TmA, in terms of their connections made in the medulla and with the LGMD in the lobula;
- the fact that one of these TmA types appears to receive direct ON input from a photoreceptor, but not the other;
- the fact that the putative ON TmA exhibits a smaller number of branching synaptic connections to the LGMD in dendritic field A, which corresponds well with the reduced intensity of ON input to field A relative to OFF input observed by Dewell et al. [16]; and
- the fact that this reduced branching would reduce the capacity for coherence selectivity mediated by reciprocal, muscarinic TmA synapses in OFF inputs [33], which is consistent with the relative lack of coherence selectivity observed by the LGMD during ON stimuli in recent experimental data [16].

While the TmA neurons in this model are purely ON or OFF selective, the choice to model them as such was not made under the presumption that they are purely so in vivo; the choice to segregate ON and OFF signals entirely was made both as a matter of simplification, and to more directly illuminate the computational consequences of differences in structure between LGMD input pathways which are known to be at least *predominantly* ON or OFF-sensitive.

One final neuroanatomical element explored here is that of potential roles for neurons presynaptic to the TmAs in the medulla. Wernitznig et al. [37] observed that of two anatomically distinct TmA types, one of them—TmA2—was not anatomically positioned to receive direct inputs from the large monopolar cells (LMCs) of the lamina, with at least one intermediate layer of neurons thus needing to exist between them. They also observed that TmA1, while positioned to receive LMC input, also received input from numerous other medulla neurons. As such, a layer of neurons has been included here between the LMCs and TmAs, providing a means to examine potential computational roles for medulla neurons in shaping the signals acting upon the TmAs.

## Methods

The model of looming-detecting circuitry in the locust presented here builds on extensive prior modelling work by Olson et al. [31], and aims to represent a complete looming detection circuit in the optic lobe of a single eye, up to and including the LGMD. It incorporates additional neurons at multiple levels within the optic lobe, and includes minor modifications to certain elements already present in the prior model. Where possible, values and structures were constrained by available electrophysiological and anatomical data, with additional values and mathematics based on detailed, well-constrained existing models. Key inferences were drawn in the process of developing the structure of the model proposed here, based on recent experimental literature and observations made in prior work [31].

### General model structure

The model encompasses several neuron types throughout the locust eye and optic lobe, including

- photoreceptors;
- large monopolar cells (LMCs);
- a hypothetical class of neurons, referred to here as intermediate medulla units (IMUs);
- a global inhibitory neuron;
- trans-medullary afferents (TmAs);
- excitatory neurons in the dorsal uncrossed bundle (DUB);
- feedforward inhibitory (FFI) neurons; and
- a single LGMD neuron The LGMD, in turn, is composed of several compartments. These include
- a series of branching compartments constituting dendritic field A, which receive excitation from the TmAs, and which converge down to a single compartment termed the distal “handle”;
- a reduced two-compartment representation of dendritic field B, receiving feedforward inhibition from ON stimuli in its distal segment;
- a reduced two-compartment representation of dendritic field C, receiving excitation from ON stimuli in its distal segment and feedforward inhibition from OFF stimuli in its proximal segment;
- a proximal “handle” compartment which couples the dendritic fields to the remainder of the neuron; and
- a spike initiation zone (SIZ) compartment, where LGMD spike generation is detected and recorded.

Fig 1 shows a reduced form of the model. This representation captures a dorsoventral slice of the model structure, a single ommatidium thick, showing the various layers of the network. The full model incorporates layers of elements arranged in a series of tessellating hexagons corresponding to individual ommatidia and their associated retinotopic units in the compound eye, similar to prior work [31]. The model diameter of 97 units across a visual field of 180° results in a total of 7,057 ommatidia and an average visual arc of roughly 1.85° per ommatidium. These values correspond well to available anatomical [42, 44] and electrophysiological [45] data.

**Fig 1.**
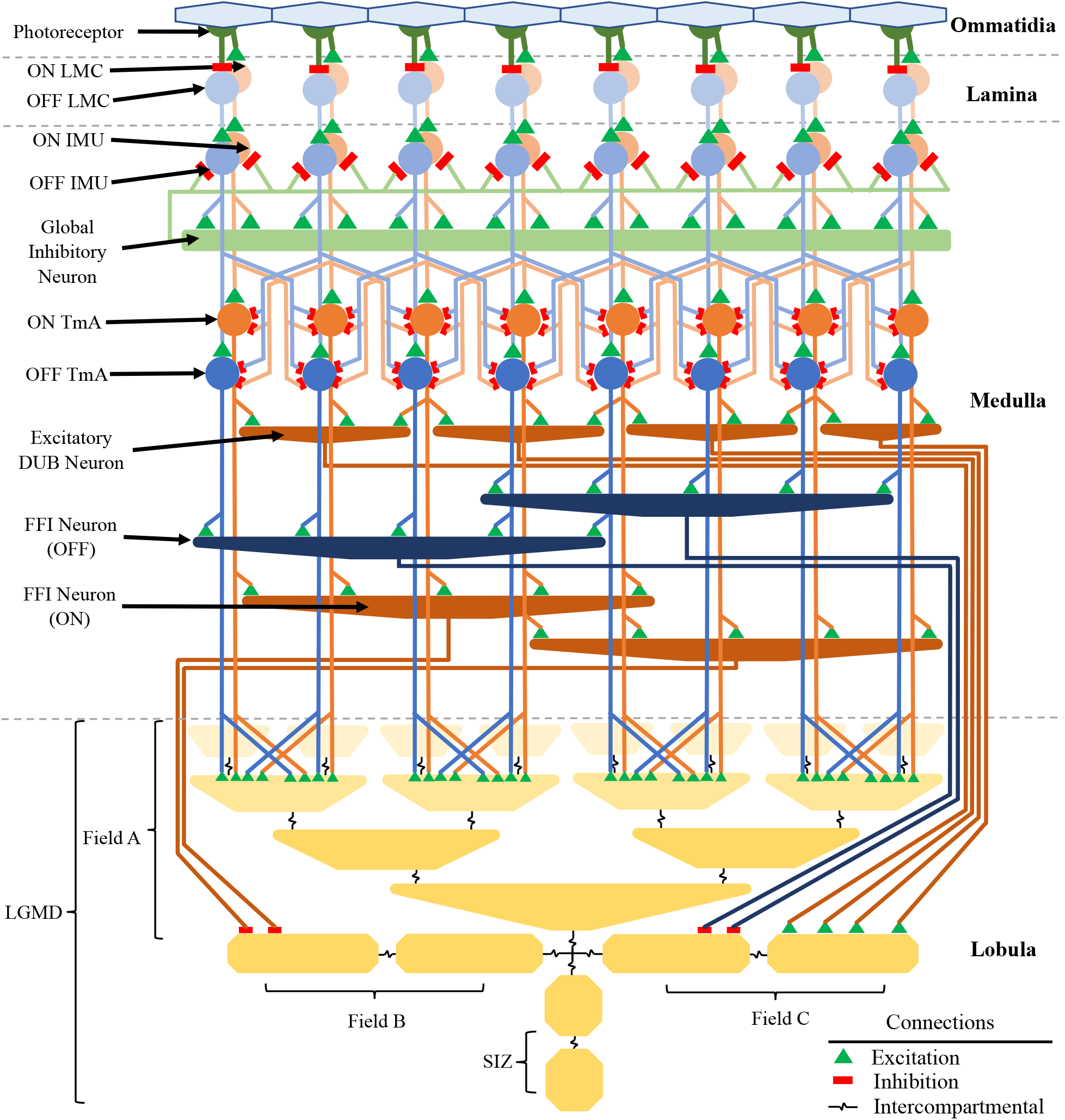
A simplified cross-section of the model structure, showing the various neuron types included, along with their layout and connections. Shades of blue and orange represent OFF and ON-sensitive neurons, respectively, with shades growing darker in the distal-to-proximal direction, while yellow is used for the compartments of the LGMD. Each of the eye’s ommatidia possesses a correspondent photoreceptor (dark green), which provides input to corresponding ON and OFF LMCs in the lamina layer. Each LMC makes excitatory connections with an intermediate medulla unit (IMU). The IMUs provide excitation to a single eye-spanning global inhibitory neuron, which in turn provides feedback inhibition onto them. The IMUs also provide excitation to their corresponding OFF or ON trans-medullary afferent (TmA), as well as providing lateral inhibition to TmAs of both their own and opposite polarities. The ON TmAs provide excitation to a smaller population of excitatory dorsal uncrossed bundle (DUB) neurons, each of which has a receptive field encompassing multiple retinotopic units. Both the ON and OFF TmAs also excite corresponding populations of feedforward inhibitory (FFI) neurons, which have wide, substantially overlapping receptive fields. Lastly, the TmAs provide excitation to Field A of the LGMD. These excitatory inputs are not constrained to their own retinotopic unit, but also branch to adjacent ones, with wider branching by the OFF TmAs (not shown). The branched structure of field A converges over the span of the eye, joining the LGMD proper alongside fields B and C, which have been simplified into two-compartment structures. The distal portion of field B and the proximal portion of field C receive inputs from the ON and OFF FFI neurons respectively, while distal field C receives input from the excitatory DUB neurons. An additional handle compartment in the LGMD couples these dendritic fields to the spike initiation zone (SIZ), which generates the LGMD neuron’s spiking outputs, and serves as the site for their detection in the model.

After the initial photoreceptor layer, most of the visual interneurons included consist of OFF/ON pairs responding to luminance decreases or increases respectively. Dark objects looming against a light background (OFF) would solely elicit responses from the former, while light objects looming against a dark background (ON) would solely elicit responses from the latter; translating objects would trigger both due to their leading and trailing edges each producing luminance transitions of opposing polarity. Separate LMCs for OFF and ON signals are consistent with current understanding of the insect visual system [46], and serve as the starting point of the separate ON and OFF pathways, which culminate in separate ON and OFF TmAs. The implementation of separate ON and OFF TmAs is based on observations of experimental data discussed earlier (see Introduction).

While the separation into OFF and ON channels in the lamina provides the computational starting point for separate OFF and ON TmAs, examination of Fig 1 shows another set of neurons included between the LMCs and TmAs, referred to as intermediate medulla units (IMUs). Unlike the LMCs and TmAs, these do not correspond directly to specific neurons in the locust. Rather, they are included for two reasons. The first is to represent the additional medulla neurons which provide inputs to the TmAs [37] (see Introduction). In particular, this resembles the input scheme for TmA2, in which at least one layer of neurons lies intermediate to the LMCs and TmAs. The second is to allow global inhibition in a feedback-type form, the possibility of which is discussed by Olson et al. [31]. This scheme was found to be required for global inhibition when testing the model’s response to looming against the background of a full-field moving grating. The feedforward structure of global inhibition used by Olson et al. [31], despite the inclusion of saturation in its value, was capable of completely shutting down TmA activity in the presence of grating backgrounds of similar contrast to simultaneous looming stimuli, which does not correspond to documented LGMD behavior [27]. On the other hand, feedback-type global inhibition is inherently self-limiting, and thus avoids completely shutting down excitatory activity while still implementing the saturation found to be required by Olson et al. [31]. Applying this feedback inhibition to the modelled LMCs directly was ineffective, as they exhibit only phasic response to inputs (which is generally true of their biological counterparts [19]). This resulted in a lack of sustained inhibition at each retinotopic unit, along with large rebound activity when the inhibition diminished.

Olson et al. [31] suggested that the TmAs provide input to additional neurons beyond the LGMD, using them as input to the inhibitory neurons of the DUB. At the time, this was justified based on the fact that the neurons of the DUB form dendritic arbours at the proximal face of the medulla [14], which the TmAs pass through en route to the lobula [36]. This possibility has since received further support from the observation by Wernitznig et al. [37] that the TmAs form excitatory synapses in the proximal medulla. This model further investigates the potential for TmAs to act as input to other neurons outside the LGMD by using them as the source of input to an additional two neuronal populations.

The first of these are the remaining neurons of the DUB, which provide excitation to field C of the LGMD that is primarily driven by ON stimuli [16]. Being constituent neurons of the DUB, they should also receive input via dendritic arbours in the proximal medulla. The feedforward inhibitory neurons of the DUB have wider receptive fields [15] than those documented by Rowell et al. [14]; it is thus likely that these excitatory neurons were specifically those DUB neurons originally observed by Rowell et al., suggesting that they receive input in the proximal medulla.

The second new group of neurons receiving input from the TmAs is a population of neurons conveying ON feedforward inhibition to field B of the LGMD. Compared to the various DUB neurons supplying input to field C, relatively little is known about these neurons in vivo. Rowell et al. [14], while lesioning various structures, were unable to remove their input to the LGMD without severing the optic chiasm containing the TmAs. They thus inferred that the inhibitory inputs to field B passed through the optic chiasm along with field A inputs. However, they did note that the component inhibitory postsynaptic potentials (IPSPs) constituting feedforward ON inhibition to the LGMD were less easily distinguished than for OFF stimuli. Based on this, the model here incorporates a greater number of ON FFI neurons than OFF, with each giving a weaker input to the LGMD.

Overall, the model presented here incorporates a number of hypothetical elements regarding the structure and function of the presynaptic network of the LGMD. While these inclusions are informed by available experimental data, they are far from verified. Consequently, this model provides an opportunity to investigate their plausibility.

As much of the work in this model is already described in detail elsewhere [31], the information presented here will focus primarily on the added element of the LGMD. However, additional minor changes have been made across the ommatidia, lamina and medulla, along with the addition of neuron types described above.

### Modifications to the ommatidia and the lamina

To better understand the effects of stimuli across the visual field of the simulated eye, alterations of the placement of ommatidia were required. In work from Olson et al. [31], the azimuthal angle of a given ommatidium was linearly proportional to its retinotopic distance from the center of the eye. This created a higher spatial density of ommatidia near the rim of the eye, as the number of ommatidia in each circumferential ring around the eye’s center increased linearly with azimuth, while the length of the circumferential ring they were placed upon would be proportional to the sine of azimuthal angle. To correct this, adjustments were made to the azimuthal spacing of the ommatidia, in order to produce a consistent density of optical axes across the eye. This had minimal impact on the response to stimuli presented by Olson et al. [31], while better allowing for the simulation of translating stimuli, as they would otherwise effectively be substantially “larger” while nearer to the edges of the visual field. It should be noted that this is a substantial simplifying assumption. Ommatidial density is not consistent across the locust eye [44], similar to the retinotopically mapped inputs to the LGMD [13, 34]. Maintaining constant spatial density of ommatidia created consistent lateral inhibitory behaviour across the eye, and simplified the creation of neurons with wide receptive fields (such as the neurons of the DUB). It also allowed clearer observation of the effects of input location versus position in LGMD dendritic field A. However, given that each ommatidial position in the model is stored as a unit vector describing its axis, future work could incorporate skewing of these axes to better reflect anatomically correct distributions.

Within the lamina, an additional population of LMCs were added to generate ON responses. These are modelled using identical equations and parameters to the OFF LMCs described by Olson et al. [31], except with a negative sign applied to their inputs. Additionally, LMC signals are no longer used as the basis for delayed lateral inhibition. Instead, this role is given to the IMUs, as described below.

### Modifications and additions to the medulla

The first addition to the medulla beyond its representation by Olson et al. [31] is that of the IMUs. Each is represented as a simple summing unit, via the equations

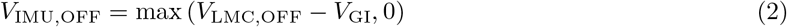

and

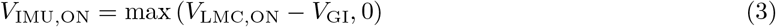

where *V*_IMU,OFF_, *V*_LMC,OFF_, and *V*_GI_ are the voltages of a given OFF IMU, its retinotopically correspondent OFF LMC, and the global inhibitory neuron, respectively [V], while *V*_IMU,ON_ and *V*_LMC,ON_ are voltages for the corresponding ON IMU and LMC [V]. The voltage of the global inhibitory neuron, in turn, is modelled by a first-order transfer function of the form

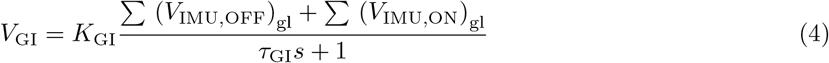

where *K*_GI_ is the global inhibitory gain, and (*V*_IMU,OFF_)_gl_ and (*V*_IMU,ON_)_gl_ are the sums of OFF and ON IMU voltages, respectively, across the eye [V]. *τ*_GI_ is the time constant governing the first-order delay applied to global inhibition [s], while *s* is the Laplace variable [1/s]. This first-order delay is a simplification, meant to encompass potential delays and elongated time constants involved in the synaptic transmission of global inhibition, along with delays in its propagation across the eye. While a more in-depth examination of global inhibition in the LGMD presynaptic circuit involving varying spatial and temporal properties warrants further study, it is not the focus of the model here.

The fact that global inhibition receives contributions from both ON and OFF channels, and applies to both in turn, was implemented based on experimental data indicating that ON and OFF stimuli are mutually inhibitory [25, 47]. The resulting inhibition should thus be highly capable at suppressing inputs from wide-field motion representing the effects of the animal’s own movement, especially in flight. Its utility in this regard is discussed later in this work.

The IMUs serve as the inputs to their corresponding TmAs, replacing the LMCs from Olson et al. [31] in this role. They also replace the LMCs as the source of lateral inhibition, although their lateral inhibitory signals are still distributed to TmAs up to six retinotopic units away, in a manner identical to that employed for the LMCs by Olson et al. [31]. In contrast with excitation, which sees OFF and ON IMUs solely connecting to their respective TmAs, each TmA receive lateral inhibition from both OFF and ON sources. This was done for the same reason as for global inhibition [25, 47]. Notably, this reduced the required lateral inhibition to suppress wide-field grating motion, and thus both the lateral inhibitory gain *K*_lat,m_ [S*/*(V · m^2^)] and time constant *τ*_lat_ [s] were adjusted substantially (see Eq 6 and 7 from Olson et al. [31] for more detail on their function). These adjusted values are presented in Table 1; other necessary values for modelling the medulla can be found in work by Olson et al. [31].

**Table 1.**
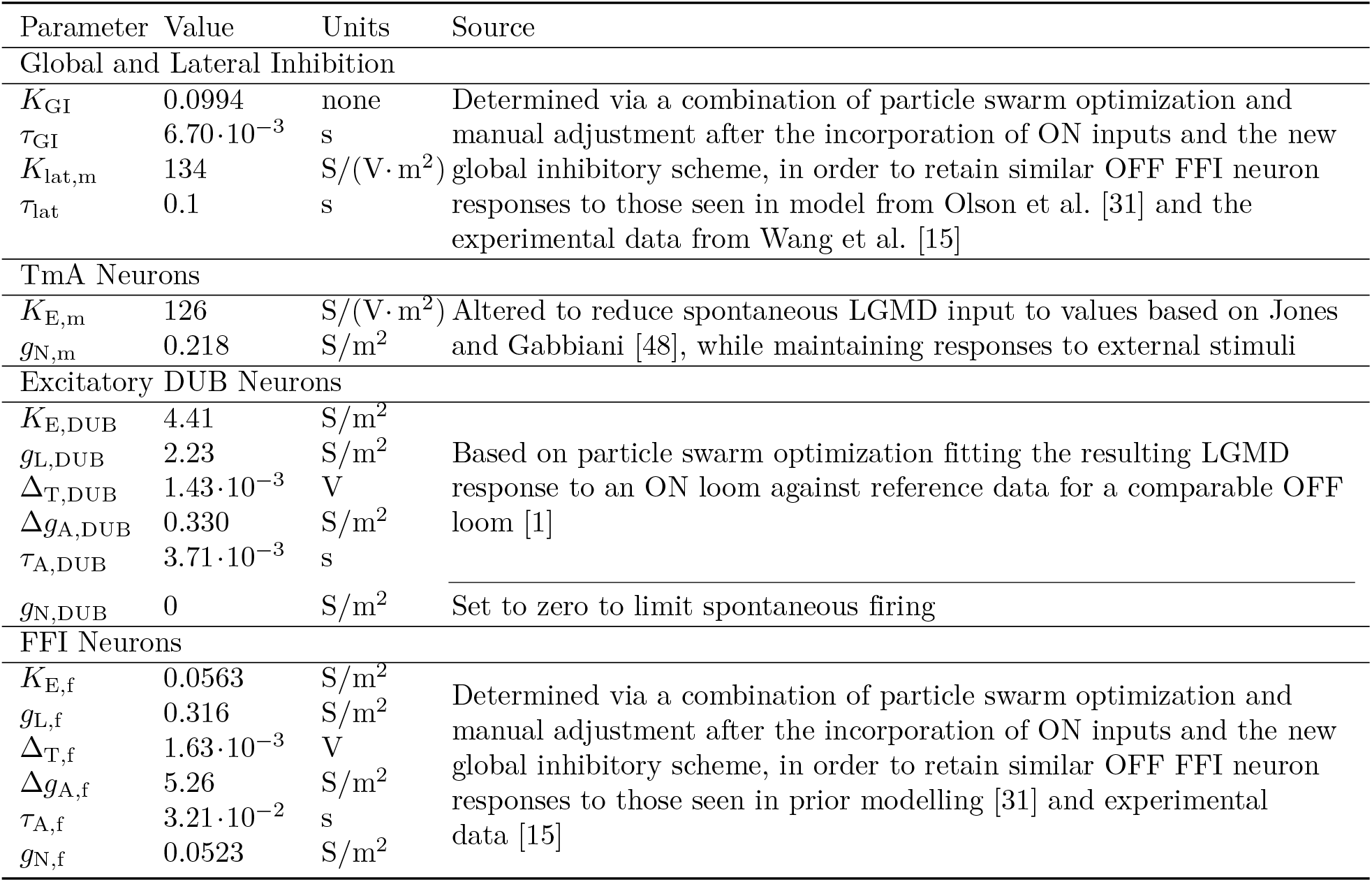
A list of parameters used in the medulla, and their sources.

Both OFF and ON TmAs are implemented with identical equations to those in the prior model. Aside from *K*_*lat,m*_, additional small changes were made to *g*_N,m_ [S*/*(m^2^)] and *K*_E,m_ [S*/*(V · m^2^)], the noise and excitatory gains. These changes were a small decrease to the former, and a small increase to the latter, reducing the frequency of randomly occurring spikes while maintaining similar stimulus response. This was done to bring the rate of randomly occurring excitatory synaptic inputs at the LGMD close to 500 Hz, which is in line with experimental observations [48]. These altered values are also presented in Table 1. While ON and OFF TmAs are given identical properties here, the relative strengths of ON and OFF inputs given to field A differ in experimental data [16]. These differences are accounted for in how their spiking generates input at the LGMD, with details provided further below (see General LGMD structure and properties).

The excitatory DUB neurons are modelled using identical equations to those used for the FFI neurons by Olson et al. [31], although with differences in parameter values and receptive field geometry. As the inhibitory neurons in the DUB have been shown to have receptive fields greater than 40° across and to constitute a minority of the DUB population [15]—an observation further reinforced by modelling work [31]—the remaining neurons of the DUB, which have been shown to carry excitation [16], are likely to be those identified by Rowell et al. [14] As such, they have been given a receptive field width of 5 retinotopic units, or approximately 9.3°, a value generally consistent with the 8°× 12° fields described for DUB neurons by Rowell et al. The receptive fields are then distributed in a similar fashion to the FFI neurons, although with a spacing between their receptive field centers of approximately 7.4°, or 4 retinotopic units. This results in a population of 469 neurons; when combined with the OFF FFI neurons, this means a total of 530 modelled neurons representing those found in the DUB, consistent with the ∼ 500 observed in locusts [14]. The parameters for these neurons (see Table 1) were determined by fitting of the response they generated from the LGMD to data for looming stimuli (see Parameter fitting for further details). Any necessary parameter values not specified were equivalent to those given for general spiking neuron parameters in Table 1 of [31]. The FFI neurons from Olson et al. [31] have been expanded to include a population responding to ON TmAs instead of OFF; these correspond to those neurons providing feedforward ON inhibition to field B of the LGMD, likely via the inner optic chiasm [14]. As few details are available about these neurons when compared with TmAs, or neurons of the DUB, much of their behavior was assumed. Namely, their receptive field sizes and internal dynamics are modelled identically to the OFF FFI neurons in the DUB. However, for reasons discussed previously (see General model structure), their receptive field spacing has been made smaller and their overall number increased relative to their OFF counterparts. This results in a receptive field spacing of approximately 7.4°, or 4 retinotopic units, and a total population of 469 neurons.

Given the changes to global and lateral inhibition, slight changes had to be made to FFI neuron properties as well. These properties were determined via similar optimization methods to those used by Olson et al. [31] (see Parameter fitting for further details); updated parameter values are given in Table 1.

### General LGMD structure and properties

The LGMD incorporated into the model encompassed a total of 133 cylindrical compartments. The main body of the LGMD consisted of three compartments: the SIZ, a proximal handle, and a distal handle (which also served as the base for field A). Fields B and C joined onto the junction of the proximal and distal handle, and consisted of two compartments each, one proximal and one distal. The remaining 126 compartments formed the branches of field A. A map of these compartments, drawn with their lengths and diameters to scale, can be seen in Fig 2.

**Fig 2.**
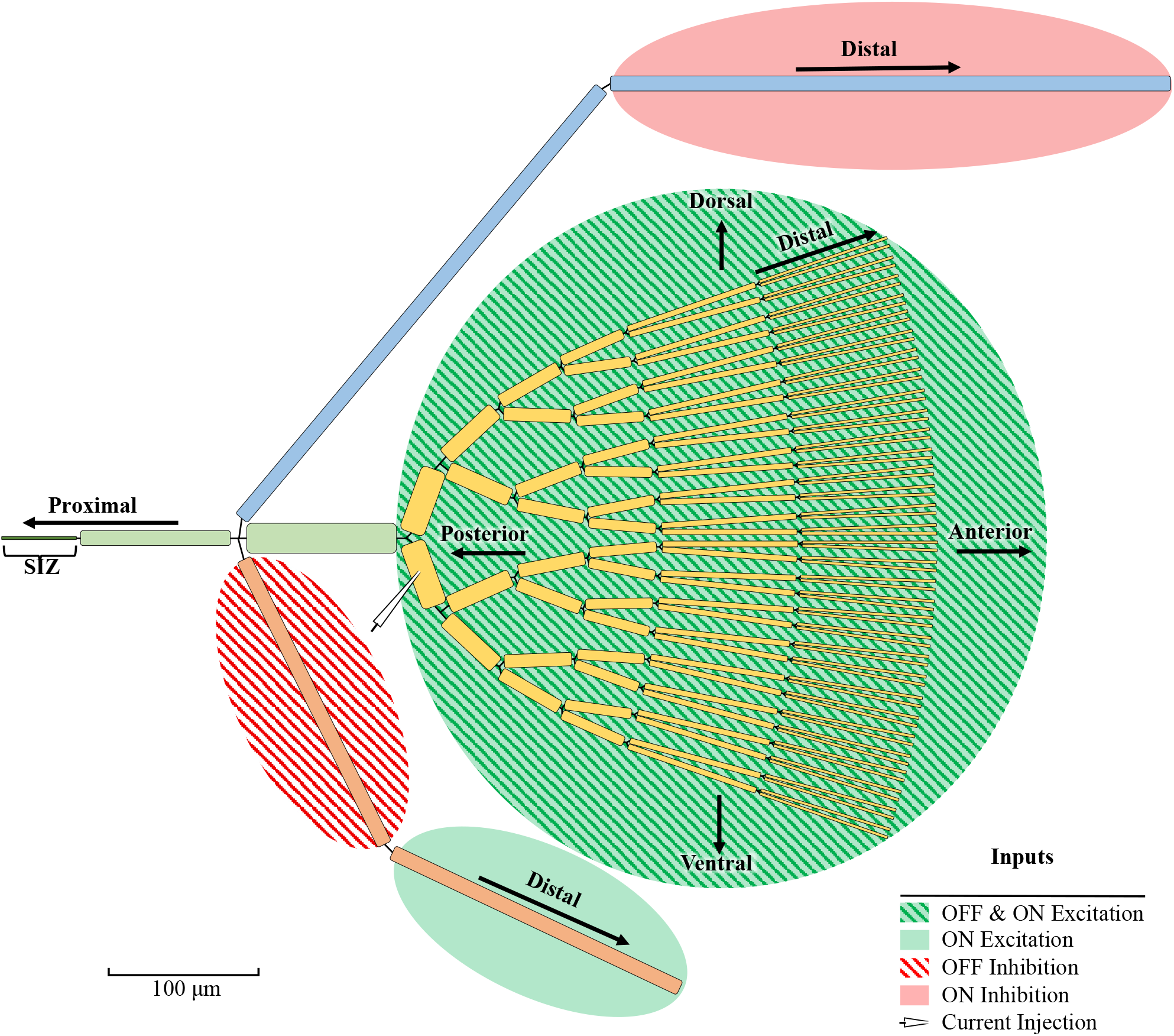
An illustration of the LGMD compartmental model, showing different functional sections and locations for inputs. Field A is illustrated in yellow, field B in blue, and field C in orange. The proximal and distal handle are illustrated in light green, with the SIZ (labelled) in dark green to the left. Proximal and distal directions along the different branching portions of the neuron are indicated with corresponding arrows. As much as possible, compartment lengths and diameters are shown to scale, with scale indicated in the bottom left. Background shaded and hatched regions indicate compartments subjected to OFF and ON excitation and inhibition. The origin of OFF and ON excitation to field A within the visual field is indicated with arrows along the dorsoventral and anteroposterior axes. The site indicated for current injections, in proximal field A, was used during determination of input resistances and membrane time constant, as well as the site for measurement of resting potential.

Where possible, LGMD geometry is informed by anatomical data, particularly those from Peron et al. [13]; further references are derived from a detailed anatomical reconstruction used in other models [16, 32, 38, 49]. Field A has a diameter of 20 µm at its base in the distal handle, tapering exponentially away from the beginning of branching towards the distal tips via the equation

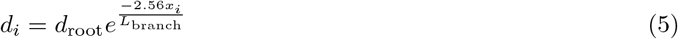

where *d*_*i*_ is the diameter of a given compartment *i* [m], *d*_root_ is the diameter at the base of field A [m], *x*_*i*_ is the location of the compartment center along the length of a given branch [m], and *L*_branch_ is the total distance from the first bifurcation in field A to a distal branch tip [m]. With a path length along each branch of *L*_branch_ = 360 µm, this results in a minimum branch diameter of 2.13 µm at the distalmost compartments.

The length of segments at each branching level is not constant, with the two distalmost branch levels having segments twice the length of the more proximal levels. This locates a greater number of visual inputs in compartments subjected to a higher degree of branching and with greater electrotonic lengths (see LGMD inputs for details on how inputs are mapped to LGMD compartments). The overall goal was to ensure sufficient electrotonic separation of inputs for spatially dependent conductances such as HCN channels and D-like potassium conductances to play a similar role in shaping LGMD behaviour as seen in vivo [32].

The overall structure used for field A (discounting the most proximal “handle” segment) possesses

- a total dendritic length of 10,090 µm;
- a mean diameter of 3.56 µm;
- a total surface area of 112,000 µm^2^; and
- a total of 126 dendritic segments.

These values are generally comparable to the average values observed by Peron et al. [13] for field A, which were

- a total dendritic length of 7,705 µm;
- a mean diameter of 3.6 µm;
- a total surface area of 108,400 µm^2^; and
- a total of 199 dendritic segments.

The most prominent difference is in the number of dendritic segments; this is likely attributable to various small, fine elements visible in dendritic field A [13, 36] being omitted in the model for simplicity.

Unlike field A, in which retinotopic mapping of inputs is both present [13, 34, 35] and computationally important [32, 33], inputs to field C appear to lack both retinotopic mapping and a computational use for it [16]. As such, modelling a large number of individual, branching compartments in field C was deemed to be a poor use of computational resources. Instead, simplifying assumptions were made about the geometry of field C in order to allow it to be collapsed into a single, electronically equivalent process, in accordance with principles laid out by Rall [50, 51]. Namely, it was assumed

- that the dendritic field exhibits branching in the form of regular bifurcation, doubling the number of branches at certain electronic distances from the field base;
- that each branch segment exhibits identical electrotonic length; and
- that at each bifurcation point, the segments had a parent:daughter diameter ratio of 2^2*/*3^ : 1.

It was also necessary to assume homogenous distribution of input across each branching level; i.e. at a given electrotonic distance from the field base. Given the non-retinotopic nature of mapping of inputs to field C [16], this would be a more reasonable assumption for field C than for the highly localized inputs to field A. It is still an approximation, but one deemed acceptable in reducing the model’s computational complexity.

With the above assumptions and conditions in place, a tree structure was generated to match the total dendritic length, mean diameter, surface area and segment count of field C as closely as possible. This dendritic tree possessed the same mean diameter and total dendritic length observed by Peron et al. [13] for field C, 1.4 µm and 2,517 µm, respectively. It also possessed a total of 127 segments, comparable to the average of 112 observed by Peron et al. This tree was then collapsed per the principles laid out by Rall [50, 51] into a single equivalent cylinder, with a diameter of 8.31 µm and length of 424 µm. This was subdivided equally into two compartments, one distal and one proximal, to allow different localization of inputs.

An identical process to the above was performed for dendritic field B, retaining the same number of segments but using a dendritic tree mean diameter of 1.6 µm and total length of 4,438 µm to match data for dendritic field B [13]. This resulted in an equivalent cylinder of 9.49 µm diameter and 748 µm length, again subdivided into a distal and proximal compartment.

The proximal and distal “handle” of the LGMD (a reference to the rake-like shape of the LGMD, used as nomenclature in other modelling work (e.g. Dewell et al. [16]) and the SIZ are each modelled as a single compartment. Compartment lengths are 100 µm for the handle sections and 50 µm for the SIZ, while diameters are 20 µm, 10 µm, and 2 µm for the distal handle, proximal handle, and SIZ, respectively. The distal handle forms the base of field A, with the equivalent cylinders representing fields B and C joined to the junction between the proximal and distal handle. This creates distances between the dendritic fields and the SIZ that are reflective of available anatomical data [13]. No axon is included in the model, with the narrow tapering of the SIZ being assumed to reduce axial current sufficiently to allow its omission. Ultimately, this does not seem to have substantially impacted the ability of the model to generate spikes or match looming responses; however, the required spiking conductance densities (i.e. those of fast sodium and delayed rectifier conductances) are somewhat higher than in some other models in the literature which incorporate a modelled axon (e.g. Dewell et al. [16]), possibly due to the absence of such conductances in an attached axon.

The general equation governing voltage in a given compartment, *V*_*i*_ [V], is

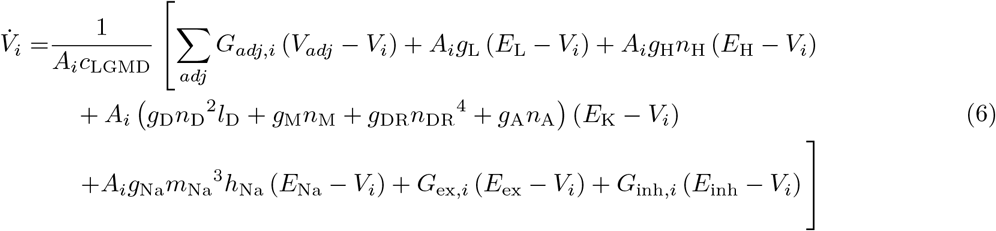

where *A*_*i*_ is the compartment surface area [m^2^] and *c*_LGMD_ is membrane capacitance [F/m^2^]. *G*_*adj,i*_ is the conductance from a given adjacent compartment to the local one [S], and *V*_*adj*_ is the voltage of said adjacent compartment. The next several terms represent various membrane conductance densities [S/m^2^], activations, inactivations, and reversal potentials [V], where

- *g*_L_ and *E*_L_ are the local leakage conductance density and reversal potential;
- *g*_H_, *n*_H_ and *E*_H_ are the conductance density, activation and reversal potential of HCN conductances;
- *g*_D_, *n*_D_^2^ and *l*_D_ are the conductance density, activation and inactivation of K_D-like_ conductances, as described by Dewell and Gabbiani [32];
- *g*_M_ and *n*_M_ are the conductance density and activation of M conductances;
- *g*_DR_ and *n*_DR_^4^ are the conductance density and activation of delayed rectifier conductances;
- *g*_A_ and *n*_A_ are the conductance density and activation of adaptation conductances, occupying the role of calcium-dependent potassium channels in the LGMD;
- *E*_K_ and *E*_Na_ are the reversal potentials for potassium and sodium conductances; and
- *g*_Na_, *m*_Na_^3^ and *h*_Na_ are the conductance density, activation and inactivation of fast sodium channels.

*G*_ex,*i*_ and *G*_inh,*i*_ are local synaptic conductances [S] for excitation and inhibition, respectively, while *E*_ex_ and *E*_inh_ are their reversal potentials. The detailed equations governing the activation and inaction variables of the various active conductances are given further below (see LGMD active conductances and spike detection).

Not all of the above membrane conductances are included in every compartment, and all vary in strength based on location even when present. HCN and D-like conductances are present only in field A (not including the proximal handle), which is where they appear to be most prominently expressed in the LGMD [32]. HCN channel densities are scaled inversely to compartment diameter (i.e. inversely to Eq 5), producing a constant per-length conductance density in the branches, and an exponentially increasing density per area from the proximal-to distalmost compartments. This is consistent with biological data, in which the voltage “sag” created by HCN channels during step current injection is more prominent at greater distances from the SIZ [32]. D-like potassium conductances possess spatially identical scaling, given their mutual participation with HCN conductances in generating coherence selectivity in field A [32].

Channels for M currents are spread throughout the entire LGMD model, but have their greatest density at the SIZ. This is due to their involvement in controlling different firing modalities [38], and is consistent with other models [16, 32, 38, 49]. Relative to their maximum strength in the SIZ, M currents have a scaling of

- 1*/*2 in the proximal handle;
- 1*/*4 in the distal handle;
- 1*/*20 in field A; and
- 1*/*80 in fields B and C.

Fast sodium channels and delayed rectifier currents are localized to the SIZ and proximal handle only, with the highest densities in the SIZ, due to their primary function being that of spike generation [52]. Initially, the proximal handle densities were 1*/*8 and 1*/*10 of those in the SIZ for the sodium and delayed rectifier conductances, respectively. These ratios were maintained while adjusting conductances for proper input resistances and resting potentials (see below and Parameter fitting for more details). Later, the proximal handle values were held constant, with the SIZ values adjusted to achieve biologically accurate responses to looming stimuli. This resulted in proximal handle densities of approximately 1*/*40 and 1*/*150 relative to SIZ values for sodium and delayed rectifier channels, respectively, as a result of large increases in SIZ conductance densities. As stated earlier, these substantial increases in firing-related conductances at the SIZ relative to other LGMD models [16, 32, 38, 49] may reflect the absence of influences from an attached axon. This is supported by the greater increase in delayed rectifier conductance than in fast sodium conductance; in the aforementioned models, delayed rectifier channels have greater concentrations relative to sodium channels in the axon than in the SIZ. Thus, the model presented here required an increase in both channel strengths—but particularly for delayed rectifiers—to achieve appropriate verisimilitude.

The adaptation current in the LGMD is generated as a result of firing, and itself serves to govern firing rate [26]. As such, it is localized solely to the SIZ and proximal handle, with the proximal handle having 1*/*3 the conductance density of the SIZ.

Leakage conductance is distributed evenly throughout the entire neuron with the exception of fields B and C, where its strength was reduced by a factor of 4. This was done to reduce electrotonic distance between feedforward inhibitory inputs and the SIZ, based on observations of LGMD behavior during the parameter fitting process. Related changes included reductions of axial resistance and relocation of feedforward inhibition (the former are discussed later in this section; for the latter, see LGMD inputs).

For intercompartmental conductances, slight changes are present as compared to how they were evaluated in prior work [31]. These changes are meant to account for the common voltage at the junction where three or more compartments meet, without directly modelling said junction, as illustrated in Fig 3. For a given compartment *i* at a junction of *n* compartments, the half-conductance *G*_*i*_ [S]—the inverse of half the compartment’s axial resistance—is evaluated as

**Fig 3.**
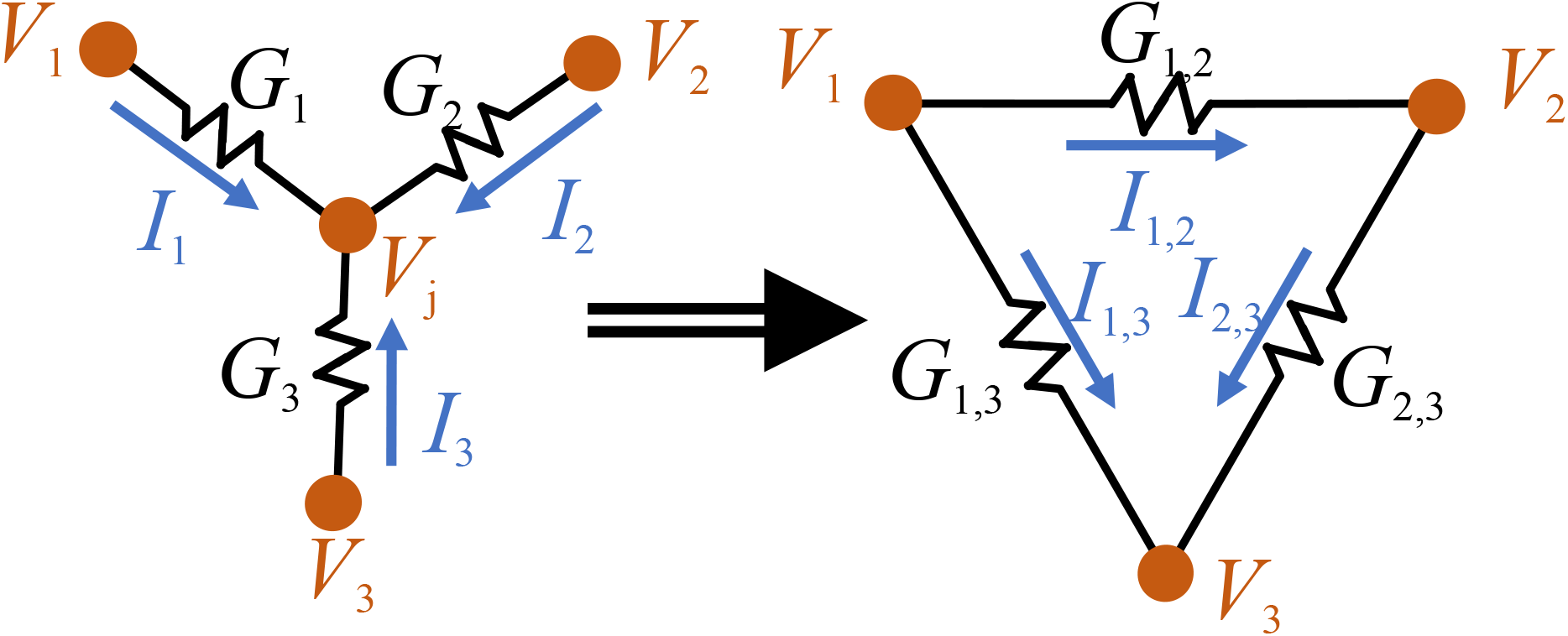
An illustration of how intercompartmental conductances were derived, for three compartments. Voltages at compartments and their shared junction are shown in orange, conductances in black, and currents in blue. The left illustrates three compartments meeting at a common junction voltage, *V*_*j*_. As the junction is a point voltage with no associated compartmental capacitance, applying Kirchoff’s current law allows a set of equivalent intercompartmental currents and conductances to be derived, of the form shown on the right.

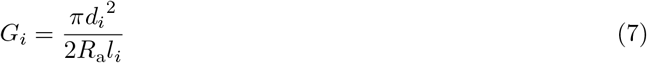

where *d*_*i*_ and *l*_*i*_ are the diameter and length of the compartment [m], and *R*_a_ is the axial resistance [Ω·m]. Then, between two compartments *a* and *b* at a junction shared by *n* compartments, the intercompartmental conductance *G*_*a,b*_ [S] is

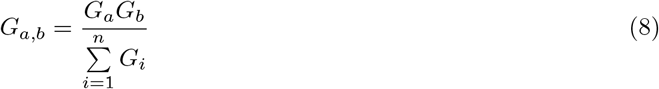

Note that for the case of only two compartments, this evaluates identically to Eq 14 from Olson et al. [31]. For instances where all but compartments *a* and *b* are at the hypothetical junction voltage *V*_*j*_ (see Fig 3), this also results in identical values for total current flow out of *a* and into *b* to the method outlined by Olson et al. (which was based on methods outlined by Dayan and Abbott [53]). However, for current flow between multiple compartments at the same junction simultaneously, Eq 7 prevents overestimation of current values, and thus underestimation of attenuation by branching structures—especially when compartmentalization of the dendritic trees is coarse. The latter point enables fewer compartments per branch segment to be used, reducing computational effort while preserving attenuation in field A.

The axial resistance *R*_a_ was estimated using purely passive membrane conductances, which were scaled along the field A branches in a similar manner to that described for HCN and D-like conductances above. Higher compartment counts were also employed in the distal segments of field A for this fitting step only, to enable inputs to be placed closer to the dendritic tip. Then, conductance and axial resistance were varied simultaneously while applying step inputs of current at the proximal handle and the distal tips of field A to achieve

- an input resistance of approximately 3.67 MΩ, measured at the first branching level of field A; and
- an attenuation of voltages generated at the distal tips of field A to approximately 1.83% of their original value when measured at the proximal handle, corresponding to an electronic distance between the two of 4 length constants, *λ*.

The former value was based on data from Dewell and Gabbiani [32, 38]. The latter was chosen based both on prior experimental and modelling work exploring the electrotonic extensiveness of the LGMD [13], and on observations of attenuation in the LGMD dendritic tree in vivo (RB Dewell, personal communication, April 7, 2025). These methods resulted in an estimated *R*_a_ of 3.3 Ω · m; this is higher than that estimated by Peron et al. [13], but similar or equal to values used in other recent LGMD compartmental models [16, 32, 38, 49]. This discrepancy with Peron et al. may be due in part to the non-uniform conductance density implemented here (and presumably in other mentioned models as well), based on observations of HCN channel density by Dewell and Gabbiani [32]. Applying this value universally across the LGMD led to excessive saturation of inhibitory inputs in field C during optimization of the looming response, however; as such, axial resistance was reduced by a factor of four in the proximal and distal handle compartments, along with the proximal compartments of fields B and C. Similar adjustments can be observed in other LGMD compartmental models [16, 32, 38, 49], suggesting that reducing attenuation in the proximal portions of the LGMD is a general requirement for modelling realistic responses (see Discussion for further comment).

After determining the axial resistance of the model, peak SIZ conductance strengths for

- fast sodium channels, *g*_Na,SIZ_;
- delayed rectifier channels, *g*_DR,SIZ_; and
- M currents, *g*_M,SIZ_

were adjusted, alongside the mean field A conductance values for

- HCN channels, *g*_H,mean_; and
- D-like potassium conductances *g*_D,mean_,

with values scaled throughout the neuron according to the ratios outlined previously. Leakage conductance and reversal potential, *g*_L_ and *E*_L_, were also varied. These values were adjusted to achieve accurate membrane potential and input resistance values for both the neuron at rest, and during blockade of HCN, M, and D-like channels, using data from Dewell and Gabbiani [32, 38] from the LGMD before and after pharmacological blockades as reference. Model input resistance was evaluated using small (0.1 nA) hyperpolarizing step currents, applied at a compartment in the first branching level of field A (see Fig 2). Once these adjustments were complete, membrane capacitance *c*_LGMD_ was adjusted to match the membrane time constant to available reference data [32, 38]. The membrane time constant was evaluated at the same compartment and using the same current inputs as input resistance, via fitting an exponential function to a window starting 2 ms after the step input, and lasting for 13.5 ms (or two time constants for the actual LGMD). This window was chosen to exclude initial influences of axial conductances, similar to the procedure used for determining time constant in work by Peron et al. [13]. The SIZ conductance density for adaptation current, *g*_A,SIZ_—more accurately described as the per-spike increment in adaptation conductance—was determined last, based on optimization of the looming response (with the SIZ strengths of sodium and delayed rectifier channels also revised at this point). This process is described in more detail further below (see Parameter fitting).

The properties for membrane conductances, axial resistances, capacitance, and reversal potentials are summarized in Table 2, along with their sources.

**Table 2.** A list of parameters governing the LGMD compartmental model, and their sources.

| Parameter | Value | Units | Source |
| --- | --- | --- | --- |
| Conductances, Capacitance and Axial Resistance |  |  |  |
| $g_{\text{L}}$ | 0.163 | S/m<sup>2</sup> | Fitting LGMD behaviour to data for input resistance and resting potentials from Dewell and Gabbiani [32, 38] |
| $g_{\text{H,mean}}$ | 1.30 | S/m<sup>2</sup> | |
| $g_{\text{D,mean}}$ | 61.1 | S/m<sup>2</sup> | |
| $g_{\text{M,SIZ}}$ | 38.5 | S/m<sup>2</sup> | |
| $g_{\text{Na,SIZ}}$ | 14900 | S/m<sup>2</sup> | Fitting LGMD looming response to firing rate data from Stott et al. [1] |
| $g_{\text{DR,SIZ}}$ | 3940 | S/m<sup>2</sup> | |
| $g_{\text{A,SIZ}}$ | 0.218 | S/m<sup>2</sup> | |
| $c_{\text{LGMD}}$ | 0.0178 | F/m<sup>2</sup> | Fitting LGMD membrane time constant to data from Dewell and Gabbiani [32, 38] |
| $R_{\text{a}}$ | 3.3 | $\Omega \cdot \text{m}$ | Chosen to achieve input attenuation in field A comparable to prior LGMD modelling work [13] and in vivo observations (RB Dewell, personal communication, April 7, 2025) |
| Reversal Potentials |  |  |  |
| $E_{\text{Na}}$ | 0.060 | V | Based on fast sodium channels other models [16, 54] |
| $E_{\text{H}}$ | -0.035 | V | Based on data for HCN conductances reported by Dewell and Gabbiani [32] |
| $E_{\text{ex}}$ | 0 | V | Based on excitatory nicotinic acetylcholine synapses in other models of the locust nervous system [18, 55] |
| $E_{\text{inh}}$ | -0.080 | V | Based on inhibitory channel reversal potentials in other LGMD models [16, 32, 38] |
| $E_{\text{K}}$ | -0.080 | V | Based on K<sup>+</sup> channel reversal potentials in other locust vision models [32, 54] |
| $E_{\text{L,lgmd}}$ | -0.0383 | V | Fitting LGMD behaviour to data for input resistance and resting potentials from Dewell and Gabbiani [32, 38] |

### LGMD active conductances and spike detection

In addition to their strengths, the voltage and time dependencies of the various LGMD membrane conductances are vital to replicating the computational properties of the neuron. Per Eq 6, most modelled channels have time-dependent gating variables governing their activation (or inactivation) within a given compartment of the LGMD model. For an arbitrary gating variable, *n*, time dependence is governed by the first-order equation

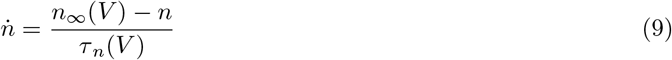

where *n*_*∞*_(*V*) is the steady-state value of the gating variable at the compartment’s current voltage, and *τ*_*n*_(*V*) is a voltage-dependent time constant [s] governing the rate of change. The exact modelling of voltage dependency for these conductances varies channel-to-channel.

For the HCN channels, the activation gating variable *n*_H_ and time constant *τ*_*H*_ are governed by equations based on those from Dewell and Gabbiani [32], which they fit to in vivo measurements of HCN channel properties. Steady-state activation gating is modelled as

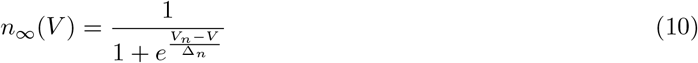

where *V* is the membrane voltage of the compartment, *V*_*n*_ is the voltage at half activation [V], and Δ_*n*_ is the activation steepness [V]. The activation time constant is given by

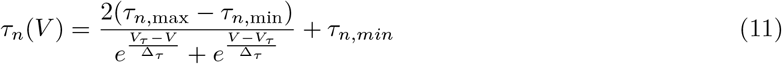

where *τ*_*n*,max_ and *τ*_*n*,min_ are the maximum and minimum time constant values [s], *V*_*τ*_ is the voltage at which maximum time constant occurs [V], and Δ_*τ*_ is the steepness of time constant change with voltage [V]. Values of the parameters in these equations are given in Table 3.

**Table 3.**
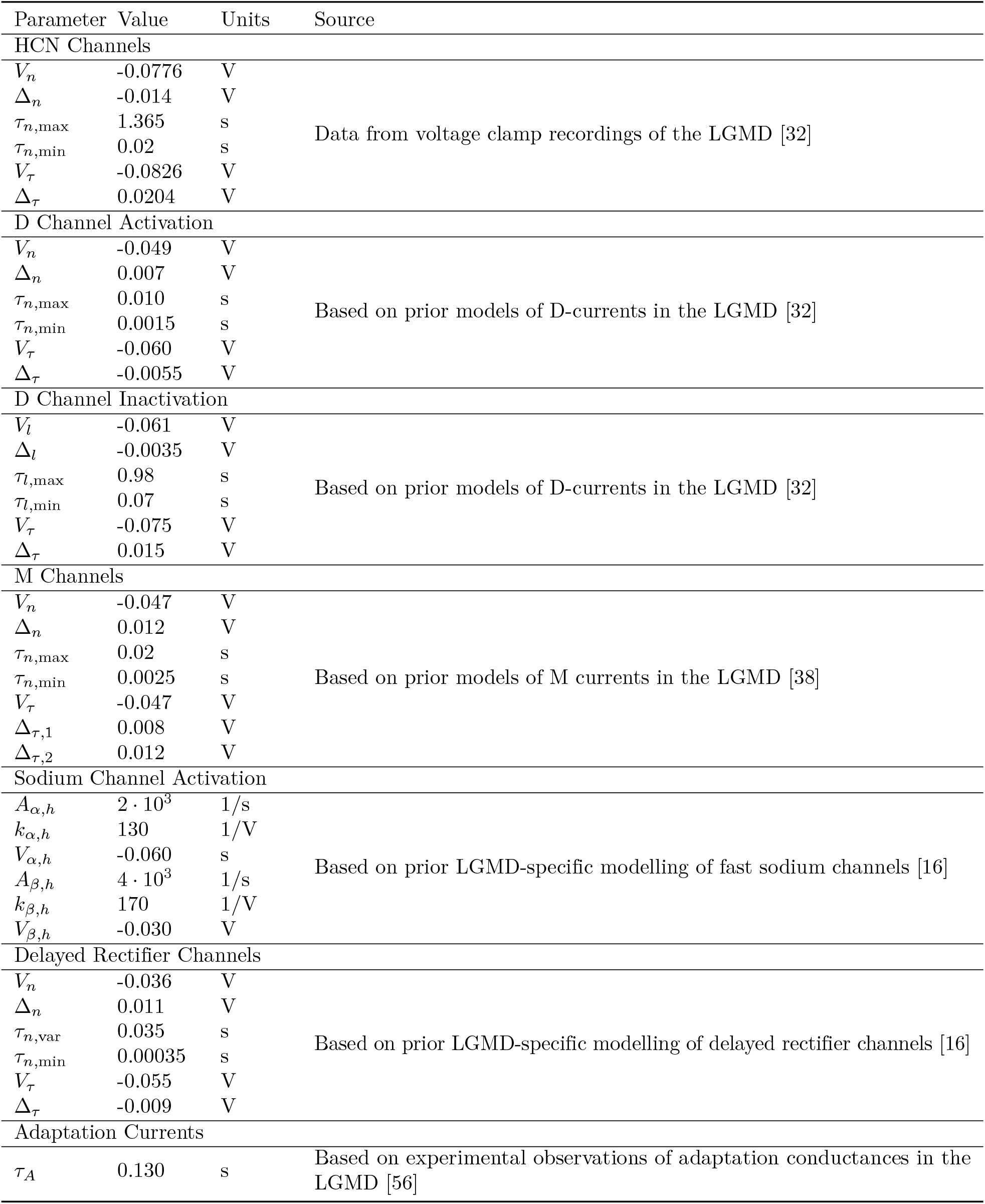
A list of parameters governing active conductances, and their sources.

The D-like conductances use a slightly different set of equations. These were derived from those used by Dewell and Gabbiani [32] to model these same conductances, in work which successfully replicated the role of D-like conductances in generating coherence selectivity in the LGMD; however, their dependence on calcium concentration was removed, as calcium was not tracked in the model presented here. The steady-state activation gating *n*_*D*_ is given by an equation of identical format to Eq 10, while time constant is modelled as

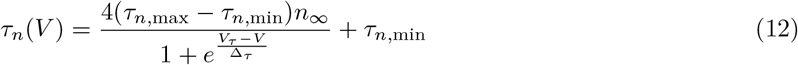

where *τ*_*n*,max_ and *τ*_*n*,min_ are the nominal maximum and minimum for *τ*_*n*_, and *n* is the steady-state gating value. *V*_*τ*_ [V] and Δ_*τ*_ [V] govern the location of maximum time constant and its change with voltage, in conjunction with the voltage-dependence of *n*. The steady-state value and time constant of the inactivation gating variable *l*_*D*_ are governed by equations of identical format to those governing the HCN channels (Eq 10 and Eq 11). The parameters governing both activation and inactivation of D-like conductances are presented in Table 3.

M currents are modelled using equations from Dewell and Gabbiani [38], as they were able to successfully capture in vivo behaviour of said channels in the LGMD. These channels have a steady-state gating variable modelled using Eq 10, identically to the other channels thus far, while the gating variable time constant was modelled using Eq 12. The parameters used in these equations for the M currents can be found in Table 3.

The modelling of the fast sodium channels is notably different from those above. The format of these equations is broadly similar to those described by Hodgkin and Huxley [52], but of a form tailored by Dewell et al. [16] to better reflect LGMD spiking. The steady-state value of the activation gating variable *m*_*Na*_ is given by

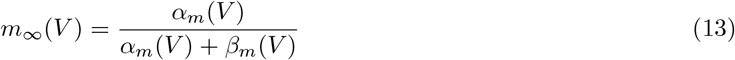

and the time constant *τ*_*m*_ is given by

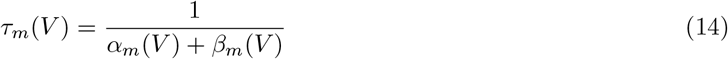

where *α*_*m*_(*V*) and *β*_*m*_(*V*) are voltage-dependent rate constants [52]. The inactivation gating variable *h* is governed by an identical set of equations. For *m*, the variables *α*_*m*_(*V*) and *β*_*m*_(*V*) are given by

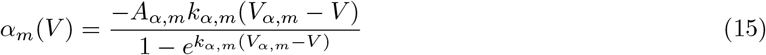

and

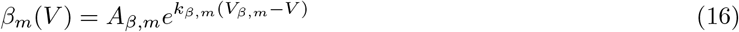

where *A*_*α,m*_ and *A*_*β,m*_ govern amplitude of the variables, *k*_*α,m*_ and *k*_*β,m*_ govern rate of change with voltage, and *V*_*α,m*_ and *V*_*β,m*_ govern voltage offset. For inactivation *h*_*Na*_, the variables *α*_*h*_(*V*) and *β*_*h*_(*V*) are given by

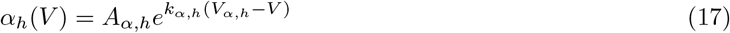

and

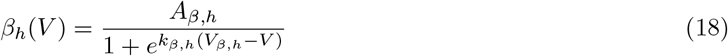

with corresponding variables serving similar roles to those in Eq 15 and Eq 16. Parameters for these equations are given in Table 3.

The delayed rectifier channels are also modelled using equations from Dewell et al. [16]. For these channels, activation gating variable *n*_*DR*_ was modelled using equations identical to Eq 10. The time constant *τ*_*DR*_ is modelled as

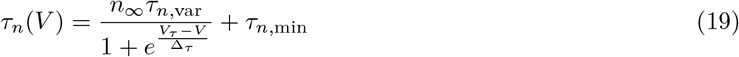

where *τ*_*n,var*_ reflects the variability of time constant with voltage, and *τ*_*n,min*_ is its minimum possible value. *V*_*τ*_ [V] and Δ_*τ*_ [V] govern the location of maximum time constant and its change with voltage, in conjunction with the voltage-dependence of *n*. The variables governing the behaviour of the delayed rectifiers are given in Table 3.

Rather than tracking calcium concentrations and fluxes as in other models [26, 54], the model here employs a simplified approach to modelling potassium currents mediating spike frequency adaptation. The activation variable, *n*_*A*_, is incremented upon detection of a spike in the SIZ such that

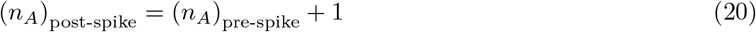

with the resulting values of *n*_*A*_ applied across all compartments where adaptation currents are present. Given that this only encompasses two tightly coupled compartments—the SIZ and the proximal handle—it was judged that spatially-varying activation could be neglected. The activation decays toward zero in a first-order fashion with a time constant of *τ*_*A*_ [s], which was given a value of 130 ms; this is consistent with the time constant of calcium extrusion in the LGMD [56]. This value is also provided in Table 3 for reference. Note that this method neglects variations in calcium influx resulting from subthreshold effects and varying action potential dynamics; in particular, this may have consequences for accurate representation of LGMD bursting properties [38].

To model the adaptation conductances as described above, it is necessary to determine when a spike has occurred. For the model presented here, the criteria were

- that SIZ membrane voltage be greater than -20 mV;
- that the derivate of SIZ membrane voltage at the previous timestep be positive; and
- that the derivative of SIZ membrane voltage at the current timestep be negative.

Spikes detected by the above method are not only used for control of adaptation conductances, but also in obtaining firing rates and spike rasters for the model.

### LGMD inputs

Setting intercompartmental conductances and various membrane channels aside, the remaining source of currents in modelled LGMD compartments is synaptic input. All modelled synapses take the form of alpha functions (see Eq. 10 from Olson et al. [31]). All excitatory synapses have a time constant of *τ*_ex_ = 0.3 ms, based on nicotinic acetylcholine synapses in other locust nervous system models [18, 55]. Inhibitory synapses have a much longer time constant of *τ*_inh_ = 12 ms; this is somewhat longer than time constants used in prior LGMD models, which have been in the range of 2-3 ms [16, 18]. This lengthening of time constant was done in response to observations made in the course of fitting the looming response of the LGMD. As inhibitory input to field C during OFF looms comes from a smaller population of neurons [15] than classically thought and is thus somewhat sparse, temporally speaking, a longer time constant was necessary to create the proper shaping of the firing rate peak and decline; shorter time constants appeared to block spiking only briefly, followed by a rebound to higher-frequency spiking (possibly due to M current deactivation during the inhibition-induced hyperpolarization). This longer time constant is still within the range of known values for decay time constants of GABA_A_ receptors [57, 58] which are known to be the source of feedforward inhibition to the LGMD [15], and thus remains biologically plausible.

For retinotopic inputs to field A of the LGMD, inputs from those ommatidia facing in the anterior direction (i.e. looking ahead of the hypothetical locust) are mapped to the distalmost portions of the dendritic tree, and those looking in the posterior direction are mapped to the most proximal compartments. This is consistent with the mapping of the LGMD in vivo [13]. The LGMD modelled here corresponds to a left-facing eye.

Mapping of excitatory synaptic inputs to field A is a multi-step process. First, at retinotopic sites corresponding to each ommatidium, inputs from adjacent facets are summed together. This reflects the branching of individual TmAs to multiple adjacent sites seen in field A [33, 35–37]. This gathering can be described as

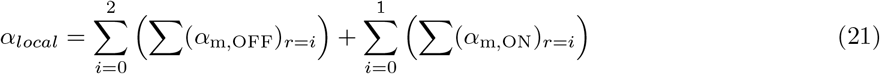

Where *α*_*local*_ is the summed input at the retinotopic site, () is the sum of all OFF TmA synaptic outputs at distance *i* from the local site, and (*α*_m,ON_)_*r*=*i*_ is the same quantity expressed for ON TmA synapses. Note that OFF inputs are gathered from a distance of up to two units away, while ON inputs are gathered from only one unit away. Moreover, the differences in the strength of ON and OFF inputs to field A come from the radius of input branching alone.

The branching distance of OFF inputs was determined based primarily on observations from Zhu and Gabbiani [35] that there was negligible overlap in activation of field A from single-facet stimuli separated by over four facets. This branching, in the model, results in inputs from a single OFF TmA arriving at 19 distinct sites, similar to the 16 synapses per TmA observed by Rind et al. [36]. The branching of the ON TmAs to sites one unit away—a total of 7 sites—was based primarily on the observation by Wernitznig et al. [37] of reduced output synapse sites in one of their two documented TmA types, which had more clear signs of ON inputs (for further discussion and justification, see General model structure). The OFF:ON ratio of outputs per TmA of 19:7 (or 2.71:1) in the model resembles the ratio of of output synapses at the LGMD of 46:15 (or 3.07:1) observed by Wernitznig et al. for their two TmA types. The resultant difference of modelled field A input strengths of 2.71:1 also strongly resembles the difference of 3.6:1 observed in calcium fluorescence observed by Dewell et al. [16]—though as they note, calcium fluorescence data is not a strict quantitative measure of input strength.

After applying both OFF and ON inputs to adjacent retinotopic sites at appropriate distances, these are mapped to corresponding branch compartments. To create this mapping, it was assumed that the number of synapses per unit length of dendrite should be held as constant as possible across field A. This assumption is based on observations by Peron et al. [13] that optic axis density at the retina varies similarly to dendritic length density of the underlying LGMD field A. However, unlike the real locust [13, 44], the model has a constant density of ommatidia across the eye. Moreover, ommatidial density and LGMD dendritic length density do not match exactly [13]. As such, this approximation should be considered an inexact one.

The means by which retinotopic input sites are subdivided and mapped to field A compartments to preserve a constant density of inputs per length of dendrite is illustrated for both a reduced retina and LGMD in Fig 4. Using this mapping scheme, the distalmost branching level of field A accounts for roughly half of the eye. At this level in field A, the branching of OFF TmAs to sites two retinotopic units away results in a single ommatidial input reaching up to four adjacent branch segments at the distalmost level. This is broadly consistent with observations of calcium fluorescence in response to single facet stimuli by Zhu and Gabbiani [35]. Once summed at a given compartment, inputs are then multiplied by a per-synapse peak conductance of *G*_A,ex_ [S] to obtain the local excitatory conductance, *G*_ex,i_ (see Eq 6). The value of *G*_A,ex_ was determined as part of fitting the LGMD response to looming data, and is given in Table 4.

**Fig 4.**
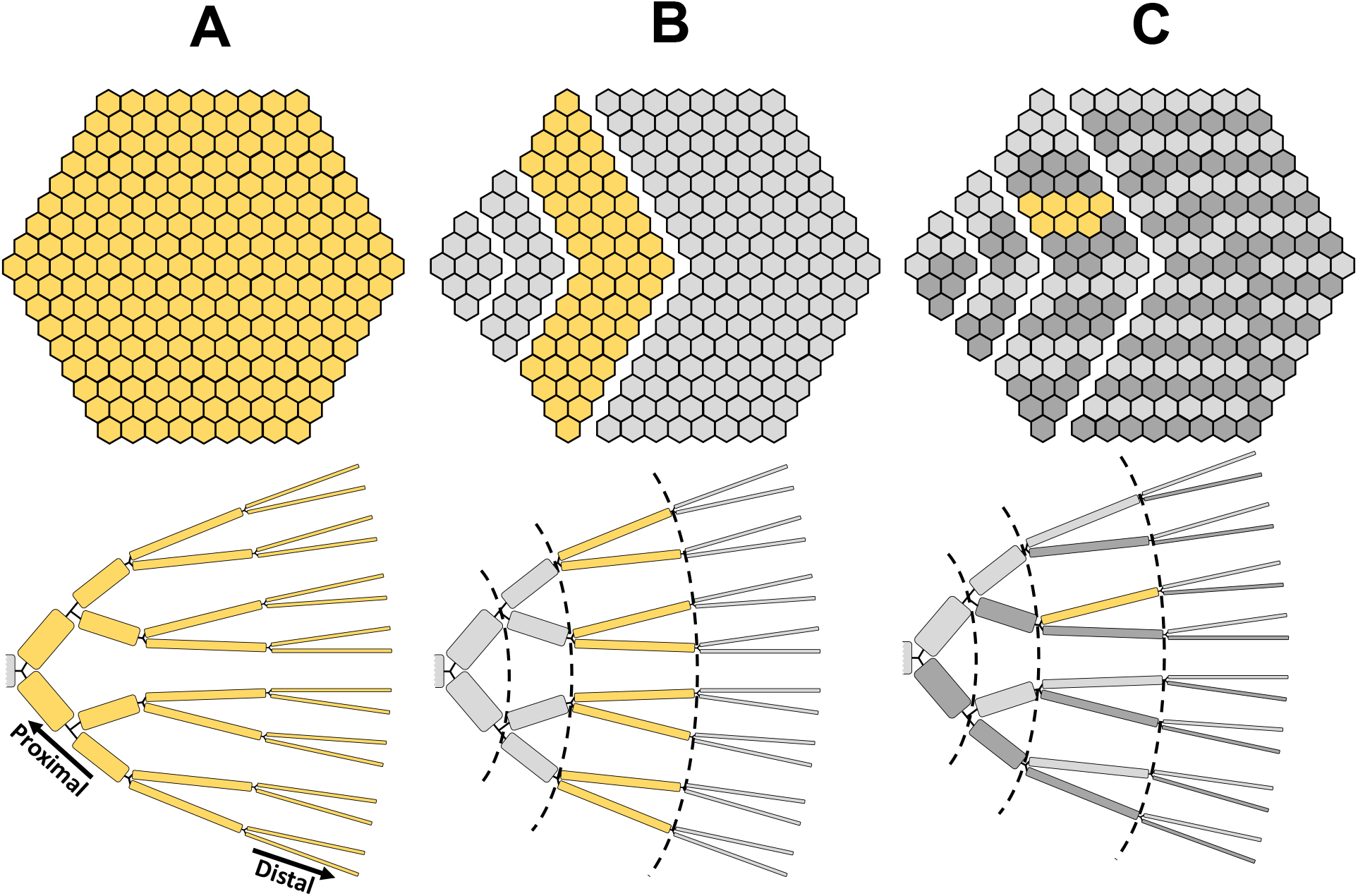
An illustration of how visual inputs were mapped retinotopically to field A, while achieving an approximately constant density of synapses per unit length. **A** shows an example of retinotopic TmA inputs and corresponding dendritic field A, reduced in number of retinotopic units and number of branching levels from the model, respectively. Note that the TmA inputs have already had branching to adjacent input sites applied, per Eq 21. **B** shows the division of the dendritic tree into different branching levels, and the division of the inputs into corresponding groups of retinotopic units at increasing distance from the leftmost point (corresponding to the most proximal portion of field A). Divisions between groups of retinotopic units are placed such that the number of units is as close to proportional to the total dendritic length present at the corresponding branching level as possible. Yellow shading indicates a corresponding field A branching level and group of retinotopic units. **C** shows the further subdivision of each unit grouping into approximately equally-sized bands mapping to different branches at the relevant branching level. An example of a single such group and its corresponding branch segment is again shown in yellow.

**Table 4.** A list of parameters governing synaptic input to the LGMD, and their sources.

| Parameter | Value | Units | Source |
| --- | --- | --- | --- |
| Synaptic Peak Conductances |  |  |  |
| $G_{A,ex}$ | $0.501 \cdot 10^{-9}$ | S | Set via fitting the LGMD response to OFF looms to data from Stott et al. [1] |
| $G_{C,inh}$ | $10.0 \cdot 10^{-9}$ | S | |
| $G_{C,ex}$ | $3.51 \cdot 10^{-9}$ | S | Scaled relative to $G_{A,ex}$ to create equivalent per-spike input conductances in field C |
| $G_{B,inh}$ | $1.30 \cdot 10^{-9}$ | S | Scaled relative to $G_{C,inh}$ to create equivalent per-loom input conductances in field B |
| Synaptic Time Constants |  |  |  |
| $\tau_{ex}$ | $0.3 \cdot 10^{-3}$ | s | Based on nicotinic acetylcholine synapses in other locust nervous system models [18, 55] |
| $\tau_{inh}$ | $12 \cdot 10^{-3}$ | s | Adjusted during optimization of the looming response to enable appropriate inhibitory function |

Outside of field A, additional ON excitation is provided by DUB neurons mapped to the distal compartment of field C. These inputs are given a per-synapse peak conductance *G*_C,ex_ that is exactly seven times *G*_A,ex_. This provides equal per-spike input strengths between ON TmAs in field A (which branch to seven different sites, per above) and excitatory DUB neurons. Once summed, these synaptic inputs provide the value of *G*_ex,i_ (see Eq 6) for the distal field C compartment. Variations in relative input strength of field A and field C inputs during parameter fitting was accomplished solely by varying DUB neuron properties to alter the number of spikes they produce. This was a simplifying assumption to limit degrees of freedom during parameter fitting, and could be better constrained in the future as more data on excitatory input to field C become available.

Inhibitory OFF inputs from the DUB are also mapped to field C. Initially, this was also done to the distal compartment; however, during fitting of the looming response the electrotonic distance from the SIZ appeared to cause these inhibitory inputs to saturate, somewhat similarly to what occurs for excitation in field A [18]. This affected the time course of inhibitory inputs reaching the proximal portions of the LGMD in such a way as to make achieving appropriate responses across varying *l/* | *v*| values difficult. It was subsequently found that moving these inputs to the proximal handle compartment improved resulting fits. In order to keep these inputs at an anatomically-appropriate location while achieving a good fit for LGMD looming responses, it was necessary to reduce attenuation via other means. As a compromise, OFF feedforward inhibition was mapped to the proximal portion of field C, and its axial resistance was reduced alongside that of the proximal and distal handle compartments (mentioned previously in General LGMD structure and properties); membrane conductances in field C were also reduced to decrease electronic distances. The peak per-synapse conductance for the inhibitory inputs, *G*_C,inh_, can be found in Table 4.

Inhibitory inputs from ON FFI neurons are mapped to the distal portion of field B, per Rowell et al. [14]. After the observations regarding field C inhibition made during the fitting of OFF looms described above, field B also received reduced membrane conductances, and reduced axial resistance in its proximal segment, though the input site itself was not changed. The strength of these inputs, *G*_B,inh_, is scaled to 61*/*469 of those in field C, based on the relative sizes of the OFF and ON FFI neuron populations. Given the otherwise identical properties of ON and OFF FFI neurons and their inputs, this should generate equivalent levels of inhibitory conductances during both OFF and ON looms, albeit with smaller and less distinguishable individual IPSPs during ON looms (consistent with in vivo observations by Rowell et al. [14]). This scaling of ON inhibition is done as a simplifying assumption given the comparatively limited data for ON looms in the literature; in the future, it may warrant further investigation.

One key element not included in the inputs to the modelled LGMD is the mutual synapses between the branching TmAs at field A [33, 36]. This was done to simplify the matter of determining TmA input strengths during coherent OFF looms. These TmA-to-TmA synapses create lateral excitation, which enhances TmA synaptic output to the LGMD when stimuli are spatially coherent [33]. As other mechanisms within the LGMD which generate coherence selectivity [32] are included in this model, the omission of this presynaptic means of creating coherence selectivity provides an opportunity to examine its relative importance. It also provides the opportunity to fit a baseline LGMD model first, making incorporating and fitting lateral excitation easier at a future date.

### Parameter fitting

Many key parameters of the model were not taken directly from literature data or reliant upon simplifying assumptions. Rather, they were adjusted via an optimization process to fit some aspect of the model’s output to experimental data for equivalent stimuli. Many of these key parameters would potentially affect the same aspects of model behaviour; as an example, both LGMD membrane conductances and synaptic strengths would affect the LGMD response to a loom. As such, a key element of the parameter fitting process was separating undetermined parameters into manageable groups that could each be constrained by comparing a different aspect of the model response to experimental data. Portions of this process have been mentioned previously, but are summarized here to provide additional details and to clarify the order of the various steps. The parameter fitting process occurred as follows:

- First, several parameters of the medulla—*K*_GI_, *τ*_GI_, *K*_lat,m_, *K*_E,f_, *g*_L,f_, Δ_T,f_, Δ*g*_A,f_, *τ*_A,f_ and *g*_A,f_—were adjusted by particle swarm optimization to match FFI neuron looming responses to data from Wang et al. [15], similar to what was performed by Olson et al. [31]. Simultaneous manual inspection and adjustment was based on comparison of responses to patches of sinusoidal grating with additional data from Wang et al. [15].
- Second, TmA parameters *K*_E,m_ and *g*_N,m_ were slightly adjusted to reduce spontaneous LGMD input to a lower frequency of ∼500 synapses per second, based on data from Jones and Gabbiani [48]. FFI looming responses were then reinspected to ensure negligible change occurred.
- A simplified version of the LGMD model was then created, with active conductances replaced by a passive leakage conductances following similar scaling throughout the neuron. The overall strength of this passive conductance was adjusted alongside *R*_a_, to determine a value of *R*_a_ that would allow an input resistance at the most proximal portion of field A of 3.67 MΩ, based on data from Dewell and Gabbiani [32, 38], while creating distal-to-proximal attenuation across field A consistent with an electrotonic distance of 4 length constants, a value similar to other models [13] and in vivo observations (RB Dewell, personal communication, April 7, 2025).
- The full LGMD model then had its input resistance and resting potential measured at the most proximal level of field A. This was done for the intact model, as well as with HCN, D-like, and M currents removed, to enable comparison with data from Dewell and Gabbiani [32, 38] for the LGMD in vivo with and without various pharmacological blockades of these currents. Conductance densities *g*_L_, *g*_H,mean_, *g*_D,mean_, *g*_M,SIZ_, *g*_Na,SIZ_, and *g*_DR,SIZ_, along with leakage potential *E*_L,LGMD_ were adjusted by particle swarm optimization to minimize errors in resting potential and input resistance under these various conditions. Within the objective function of the particle swarm optimization, double weighting was given to errors in the resting potential and input resistance of the intact model, to prioritize its correct behaviour.
- The LGMD model then had *c*_LGMD_ adjusted to achieve a membrane time constant at the most proximal level of field A of 6.76 ms, based on data from Dewell and Gabbiani [32, 38].
- The LGMD model was connected to the OFF and ON TmAs and FFI neurons, and the completed model subjected to OFF looms of discs at *l/* | *v* | = 23 ms and 93 ms. Parameters *G*_A,ex_, *G*_C,inh_, and *g*_A,SIZ_ were adjusted via particle swarm optimization to fit the modelled LGMD firing rate to DCMD firing rate data for equivalent stimuli from Stott et al. [1]. *g*_Na,SIZ_ and *g*_DR,SIZ_ were also adjusted, albeit with the effects limited to the SIZ compartment only. Error in firing rate for each *l/* | *v*| was calculated over a window starting 1 s prior to collision and ending 0.125 s after, with the sum of squared residuals between the model and experimental data normalized to the summed experimental firing rate data over the same window. The objective function value used in the particle swarm optimization was a weighted average of this error at the two specified *l/*|*v*| values, with the weighting being the error values themselves. Thus, the *l/*|*v*| with higher error predominated the objective function value, to discourage the optimization from over-fitting to data for a single *l/*|*v*|.
- Lastly, the excitatory DUB neuron parameters *K*_E,DUB_, *g*_L,DUB_, Δ_T,DUB_, Δ*g*_A,DUB_ and *τ*_A,DUB_ were adjusted via particle swarm optimization, based on comparison of the LGMD firing rate response to ON looms at *l/* |*v*| = 47 ms with experimental data from Stott et al. [1] for OFF looms at the same *l/* |*v*|. The baseline firing rate of DUB neurons was also assessed, and used to apply a quadratically-scaling multiplicative penalty to the objective function value for the particle swarm optimization at cumulative firing rates greater than 1 Hz.

For particle swarm optimizations, convergence was determined to have been achieved when the relative change in the objective function optimum was less than 10^*−*6^ over a duration of 20 iterations.

A few things should be noted about the above process. The first is that—aside from the parameters of excitatory DUB neurons and minor adjustments to spontaneous TmA firing—the presynaptic network feeding the LGMD was only adjusted on the basis of feedforward inhibitory neuron firing in the DUB, similar to what was done by Olson et al. [31]. Fitting of the OFF loom response of the LGMD was done solely by adjustment of the LGMD itself.

The second point to note is that the ON looming response of the LGMD was fit to experimental DCMD data for OFF looms. In the literature there are few firing rate data for complete responses ON looms, and none directly comparable to the OFF looming data used to fit the LGMD response. However, Dewell et al. [16] observed no significant difference in jump timing between comparable ON and OFF looms at *l/* |*v*| = 40 ms. Jump timing is directly correlated with the peak and subsequent decay of LGMD firing rate [5], suggesting that firing rate peak times should be roughly equivalent at similar *l/* |*v*| values. Moreover, the LGMD firing rate magnitudes documented by Dewell et al. [16] were also similar between coherent ON and OFF looms—although they highlight that these were from different animals, and thus direct quantitative comparisons are of limited validity. Mitra et al. [59] presented direct comparisons of the timing and magnitude of peak firing rate between ON and OFF looms across multiple *l/* |*v*| values. Their data show minimal difference in peak time between ON and OFF looms at *l/* |*v*| = 50 ms for *Schistocerca americana*, although the difference for *Schistocerca gregaria* at *l/* |*v*| = 40 ms is more noticeable. Nonetheless, OFF looming data from Stott et al. [1] for *l/* |*v*| = 47 ms permitted fitting of qualitatively correct ON looming responses—especially when compared to the model’s OFF responses—even if not necessarily producing quantitatively exact ones.

### Simulations and stimuli

The model described above was simulated using MATLAB Simulink. The various non-spiking components were simulated using an Euler ODE solver with a step size of 0.4 ms as per Olson et al. [31], while spiking medulla neurons were simulated using an Euler ODE solver with a step size of 0.05 ms, as this produced results functionally identical to the 0.01 ms step size used by Olson et al. while requiring far less computation time. The LGMD compartmental model was simulated using the ode45 variable-step solver within Simulink, with a relative tolerance of 10^*−*6^ specified. The solver was chosen from the available variable-step solvers in Simulink, which included both stiff and non-stiff options. Selection of the solver and tolerance was performed on the basis of being able to produce analytically correct simulations of simplified passive multicompartmental models and avoid spurious oscillations of membrane voltage in a full-scale simulation of a passive field A, while requiring the minimum computational time to do so.

The completed model incorporates a large number of states, totalling

- 28,229 states across all non-spiking neurons (photoreceptors, LMCs, and simulations of lateral and global inhibition);
- 60,452 states across all spiking neurons in the medulla and DUB; and
- 651 states in the LGMD model.

Simulations were performed on a computer equipped with an 8 core Intel Core i7-11700 processor, running at 2.50 GHz. Simulations of neurons presynaptic to the LGMD took approximately 330 s to run per second of simulated time, while simulating the LGMD took approximately 40 s per second of simulated time, with up to eight simulations being run in parallel at a given time.

For each given stimulus, 19 separate runs of the model were performed. For each of these runs, two factors were varied:

- the center of the OFF FFI neuron population was shifted to various retinotopic unit offsets relative to the center of the retina, per Fig 7 of Olson et al. [31]; and
- the central axis of the eye was offset in the anterior (forward) direction from the hypothetical lateral axis of the animal in equal steps from 0 to 1.7 degrees.

As with the offsets in stimuli used by Olson et al. [31], this compensated for the influence of the highly regular and symmetric spacing of neurons in the model on the relative timing of TmA and OFF FFI neuron firing. This variation of neuron positioning relative to stimuli also created inter-run variability analogous to inter-trial and inter-animal variability in reference experimental data.

To verify comparable FFI neuron behaviour to that seen in Olson et al. [31], the model up to and including the medulla was subjected to OFF looms of discs approaching the center of the eye, with *l/* |*v*| of 20, 40 and 80 ms. As the offsetting of OFF FFI neurons performed here is geometrically identical to the offsetting of the stimulus relative to the neurons performed by Olson et al., the outputs of the two models can be directly compared. These stimuli also permit comparison with experimental data for equivalent stimuli from Wang et al. [15].

To examine the model’s ability to reproduce the qualitative and quantitative aspects of LGMD firing under a variety of conditions, diverse stimuli were employed. These are illustrated in Fig 5, and consisted of

- OFF looms of discs approaching the center of the eye (Fig 5A), with *l/* |*v*| of 23, 47 and 93 ms, directly comparable to experimental data from Stott et al. [1];
- similar ON looms (Fig 5A) at *l/*|*v*| = 47 ms only;
- incoherent ON and OFF looms (Fig 5B) at *l/* |*v*| = 47 ms with ommatidial luminance values shuffled to random sites elsewhere on the retina for each trial, resembling (though not identical to) to methods used experimentally by Zhu et al. [33], Dewell and Gabbiani [32], and Dewell et al. [16] to create incoherent looming stimuli;
- A black disc looming with *l/* |*v*| = 47 ms, presented against a background of a sinusoidal black-and-white grating occupying the entire visual field (Fig 5C), with a grating period of 13.3° and angular velocity of 28.4°/s;
- The above stimulus repeated against a blank background for comparison (Fig 5C); and
- a 3.5 cm radius disc moving at 3 m/s through a point 80 cm away along the central axis of the eye, with motion consisting of looming (*l/* |*v*| = 12 ms), anterior-to-posterior translation, or transitions from translation to looming or looming to translation at the point (Fig 5D).

**Fig 5.**
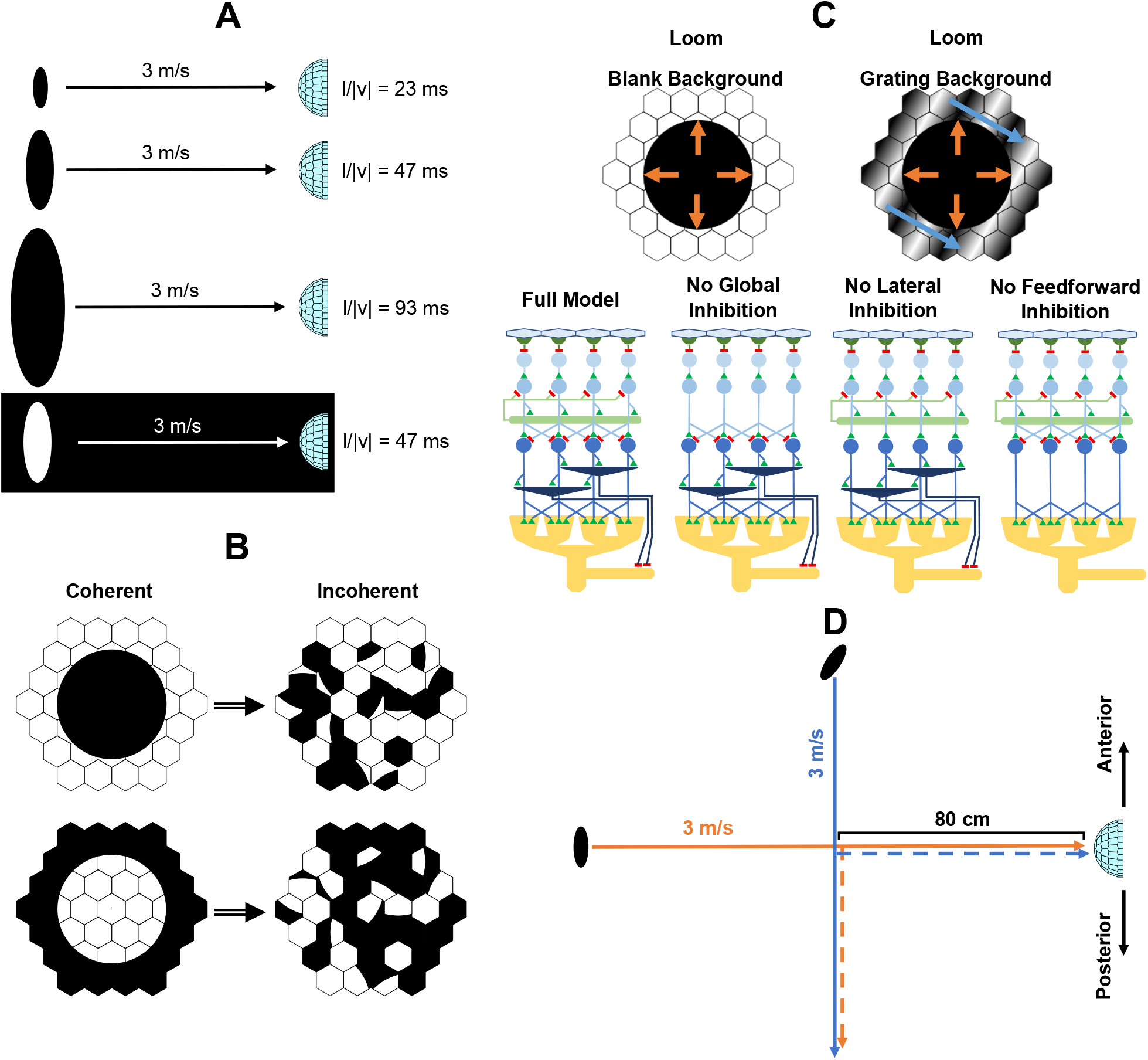
An illustration of various stimuli and configurations used to test the LGMD model. **A** shows basic OFF looming stimuli consisting of a black disc against a white background, approaching the simulated eye along its central axis. These stimuli vary the disc radius, *l*, to create a range of *l/* | *v*| values. Also included is a single ON loom, consisting of a white disc against a black background. **B** shows representations of how incoherent OFF and ON looms were created from their standard counterparts, on an example retina with reduced ommatidia count. Each ommatidial luminance input was shuffled to another random site, creating an image of equal total luminance, but lacking spatial coherence. **C** shows a similarly reduced representation of an OFF loom against a blank background and that of a moving sinusoidal grating. For these stimuli, grating movement started first, followed by object expansion. Also shown are representations of the various model structures tested against these two stimuli; for simplicity, only the OFF-sensitive neurons are shown, with representations matching the color scheme of Fig 1. The model configurations tested included the full model, as well as models with global, lateral, or feedforward inhibitory elements removed. **D** shows various stimuli combining translatory and looming elements. All stimuli, consisting of a 3.5 cm radius black disc, passed through a point 80 cm away on the eye’s central axis at a constant speed of 3 m/s. The disc remained perpendicular to the eye at all points. Orange trajectories reflect those that were initially looming, while blue trajectories reflect initial translation. Dashed lines show trajectories that changed direction. Translation started at a point anterior to the modelled eye, and proceeded in the posterior direction.

All looming stimuli stopped a single timestep before colliding with the eye, resulting in a final subtense angle of almost 180°.

For incoherent stimuli, the random mapping used in their generation was changed in each of the 19 model runs performed. A fixed seed was used for the random number generator such that the set of 19 mappings was identical for all incoherent stimuli, to permit more direct comparisons between them.

For the stimuli presented against a grating background, grating motion was initiated 1.5 s after the start of the simulation, at a time 3 s prior to the collision of the disc with the eye, with the initiation of looming occurring 2 s prior to collision. This corresponds to an initial object subtense angle of 2.69°, close to the arc surveyed by a single modelled ommatidium. The blank background stimulus used for comparision had an identical start time for looming. Unlike other stimuli, trials for looming in the presence of a grating background were run using not only the intact model, but also with the following elements removed:

- global inhibition in the medulla;
- lateral inhibition in the medulla; and
- feedforward inhibition to the LGMD.

These four configurations were also tested for the matched looming stimulus without a grating background, in order to establish the functions of different inhibition types during looming in environments both with and without background visual input.

The last set of stimuli, incorporating combinations of translation and looming through a single common point, were designed to be directly analogous to stimuli used by McMillan and Gray [23]; this is true for the looming, translating, and translation-to-looming stimuli. A stimulus which transitioned from looming to translation was also included, matched to the trajectories exhibited by the pure looming and translating stimuli. No direct analogue to this exists in the experimental data, though other stimuli featuring transitions from looming to translating motion are present in work by McMillan and Gray [23] and Santa Rita et al. [60] which allow qualitative comparison.

The incoherent stimuli mentioned above produced results which warranted investigation via further alterations to the model and stimulus; these will be described later (see Results) as they become relevant.

## Results

While the primary focus of this manuscript is on the responses of the LGMD, the changes made to the ommatidial axes and to the modelling of the medulla requires re-examination of the behaviour of the feedforward inhibitory neurons of the DUB to ensure that they accurately capture in vivo behaviour. Fig 6 shows that looming responses from the OFF FFI neurons are consistent with their counterparts work from Olson et al. [31] The results suggest that implementing global inhibition in a feedback form, as permitted by inclusion of the IMUs, is capable of replacing the “saturation” implemented via Eq 8 of Olson et al. [31]. Additionally, the strength of lateral inhibition is reduced compared to [31]. Their work suggested that global inhibition is more important than lateral inhibition in shaping the response of OFF FFI neurons to looming, with the importance of lateral inhibition lying elsewhere. The new model reinforces this point by replicating biological looming responses with reduced lateral inhibition.

**Fig 6.**
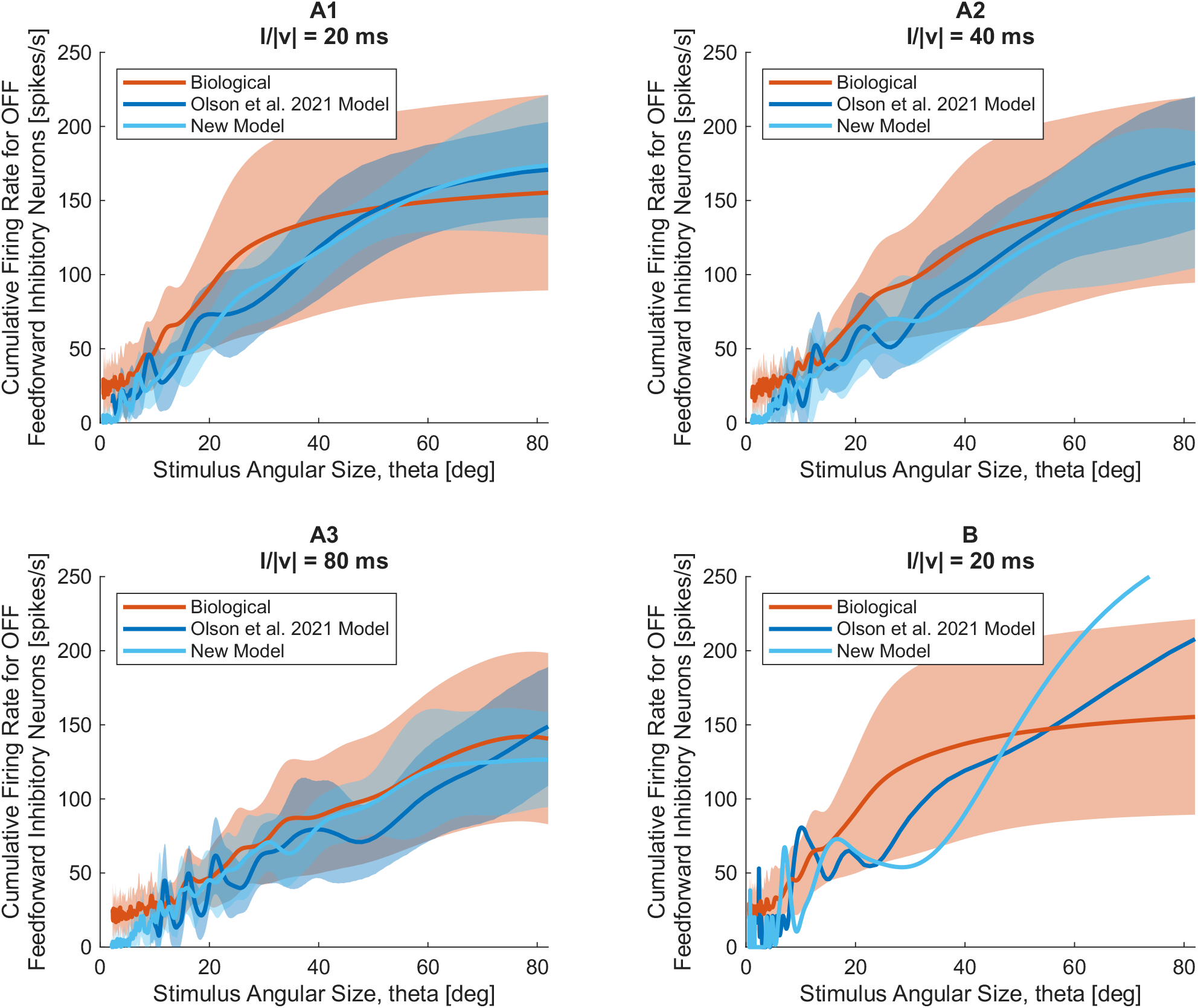
Responses of modelled FFI neurons in the DUB to looming stimuli, compared with prior modelling work from Olson et al. [31] and biological data from Wang et al. [15]. Model firing rates were summed across the entire population of simulated FFI neurons, and were obtained from a spike histogram with 1 ms bin width, smoothed by a Gaussian convolution with standard deviation of 20 ms; biological data were processed using an identical Gaussian [15]. **A1-A3** show responses for stimuli with *l/* | *v*| of 20, 40, and 80 ms, with data from both models averaged across multiple model runs (*n* = 19). Each run employed a different offset of the FFI neuron receptive fields from the line of stimulus approach. Data from Wang et al. [15], averaged across multiple animals (*n* = 6), are plotted for comparison. Shaded regions indicate the standard deviation of model outputs across trials (light blue for the current model, dark blue for Olson et al. [31]) and of experimental data across animals (orange). **B** shows representative single runs from each model at *l/*|*v*| = 20 ms, with the stimulus approach centered on the map of FFI neurons.

### Simple looming responses

Following verification of FFI behaviour, it is logical to examine the modelled LGMD responses to looming (the preferred stimulus of its biological counterpart). Fig 7 shows a variety of model loom responses, with experimental data from Stott et al. [1]. Modelled firing rates in response to OFF looms are generally consistent with biological data across tested *l/* |*v*| values. This is expected for *l/* |*v*| = 23 ms and 93 ms, as these were used directly in fitting model response. *l/* |*v*| = 47 ms, however, was not, and thus provides a small test of the model’s predictive abilities. On a qualitative level, the model performs well, with the 47 ms response having a firing peak time and magnitude somewhere between the 23 ms and 93 ms, similar to biological data.

**Fig 7.**
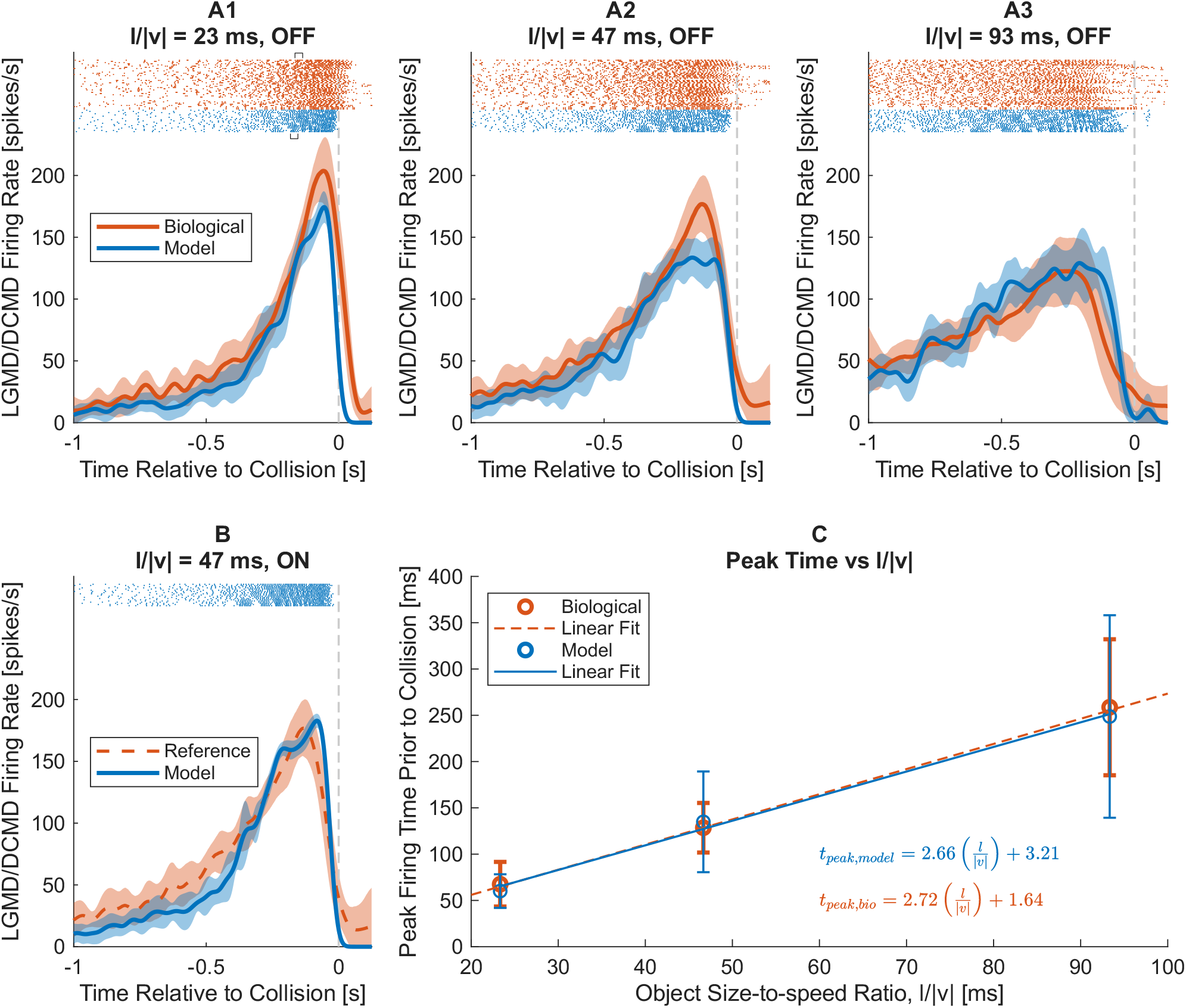
Responses of the modelled LGMD to looming stimuli, compared with biological data from Stott et al. [1]. Model firing rates were obtained from a spike histogram with 1 ms bin width, smoothed by a Gaussian convolution with standard deviation of 21 ms, for comparability to the 50 ms full width at half maximum (FWHM) Gaussian used by Stott et al. [1]. Model firing rates were averaged across multiple model runs (*n* = 19), while experimental values were averaged across multiple animals (*n* = 14, 3 trials per animal). Dashed vertical gray lines indicate the time of collision. **A1-A3** show firing rates for OFF looming stimuli with *l/* | *v* | of 23, 47 and 93 ms. Shaded regions indicate standard deviations of model output (across runs) and experimental data (across animals). Spike rasters for all 42 experimental trials and 19 model runs are shown above. Examples of high-frequency clustered spikes are indicated on the top experimental spike raster and bottom model spike raster of **A1** by black brackets. **B** shows the model firing rate for an ON looming stimulus with *l/* | *v*| = 47 ms, with data from Stott et al. [1] for an otherwise identical OFF loom plotted for reference. Shaded regions represent standard deviations as above, and spike rasters for the 19 model runs are shown at the top. **C** shows a plot of the time of of peak firing rate relative to collision versus *l/* | *v*| for both model and experimental data. Bars indicate standard deviations of model (across runs) and experimental (across animals) peak times. Linear fits to peak time versus *l/* | *v*| are shown for both the model (*R*^2^ = 0.552) and biological reference data (*R*^2^ = 0.849), along with their equations. Data from Stott et al. [1] used to create the linear fit included additional points at *l/*|*v*| = 13 ms and 140 ms (not shown).

While the overall trajectory of firing rate is similar between the model and biological data in all three cases, there are noteworthy differences. Specifically, the peak of the average model firing rate is broader than its experimental counterpart, especially for higher *l/*| *v*| values. Model standard deviation around the peak time also grows larger as *l/*| *v*| increases. This suggests that the broader firing rate peak is not purely a phenomenon inherent to every model run, but rather appears in the average as a result of greater inter-run variability in peak time at higher *l/* |*v*|. Although the variability in modelled peak time is larger that for biological data (Fig 7C), it is still within a similar order of magnitude to that seen biologically.

Examination of spike rasters yields insights into the source of peak time variability in the model. The model responses exhibit spike clusters (see Fig 7A1) which resemble burst firing that characterizes DCMD (and thus LGMD) firing during the latter part of a loom [39]. This clustering becomes more apparent in the model outputs across a wider temporal range as *l/* |*v*| increases. These “bursts” vary both in their positioning, and local spike rate from run to run, with the corresponding variation in firing rate on short time scales creating highly variable peak times. Within the model, only two factors were directly varied on a run-to-run basis:

- the positioning of the OFF FFI neurons; and
- the slight angular offset of the stimulus approach.

As such, one or both of these elements likely create the variation in model bursting—and peak time—seen at higher *l/* |*v*|.

Examining the model’s ON loom response, seen in Fig 7B, also yields a generally close fit to the reference data, but also a few noteworthy points of difference from the model’s OFF responses. As compared to the 47 ms OFF loom, the ON stimulus generates a higher peak firing rate, closer to that seen in the reference data. Given that this reference data is for an OFF loom, such similarity is not directly meaningful for the model’s veracity; in fact, a distinctly higher peak firing rate in response to an ON loom would be unlikely in vivo [12, 16].

While assessment of the looming response thus far has generally been qualitative, one easily quantified aspect is the time of peak firing rate. A linear increase in peak time prior to collision with increasing *l/* |*v*| for looming stimuli is a key characteristic of LGMD behaviour [1, 20, 61, 62], likely reflecting the way in which field A excitation and feedforward inhibition interact in the neuron [2, 18, 20]. The model reproduces this extremely faithfully, with close similarity in mean peak times between the model and experimental data from Stott et al. [1] visible in Fig 7C. The seemingly low *R*^2^ of the linear fit to model data, relative to the linear fit to experimental results from Stott et al., is attributable to two factors. The first is that the experimental data encompasses a wider array of *l/*| *v*| ; calculating the experimental *R*^2^ based only on data from *l/* |*v* |values also examined in the model causes its value to drop to 0.747. The second factor limiting the comparability of the two *R*^2^ values is the greater variability in peak time in the model response, as discussed previously. However, even for LGMD and DCMD firing observed in vivo [1, 17], it is the *mean* time of peak firing which shows a strong linear dependence on *l/* |*v* |, with variability between trials and animals still being apparent; as seen in Fig 7C, the standard deviations of peak time observed by Stott et al., while lesser than those seen in the model, are still of a similar order of magnitude. When considering the mean values of peak time alone, the linear fits to model and experimental data achieve *R*^2^ values of 0.996 and 1.000, respectively. This illustrates not only the extreme linearity of the relationship between mean peak time and *l/* |*v*| in the LGMD, but also the model’s extremely accurate replication of this property. Thus, while the model reproduces basic looming responses in a way that is qualitatively faithful, it also reproduces key quantitative elements of LGMD behaviour with high fidelity.

### Stimulus coherence

While the ability of the model to reproduce looming responses is of interest, an actual LGMD neuron must be able to deal with more complex stimuli—and so too must an effective model. The first such stimulus presented to the model was that of incoherent looms, the responses to which can be seen in Fig 8. While the actual LGMD neuron responds preferentially to coherent OFF looms over incoherent ones [32, 33], the model shows an inverse preference (Fig 7A), with a generally greater response to incoherent looming stimuli. The preference for incoherent stimuli becomes particularly pronounced after ∼250 ms prior to collision, resulting in 48% increase in spiking overall relative to the coherent stimulus.

**Fig 8.**
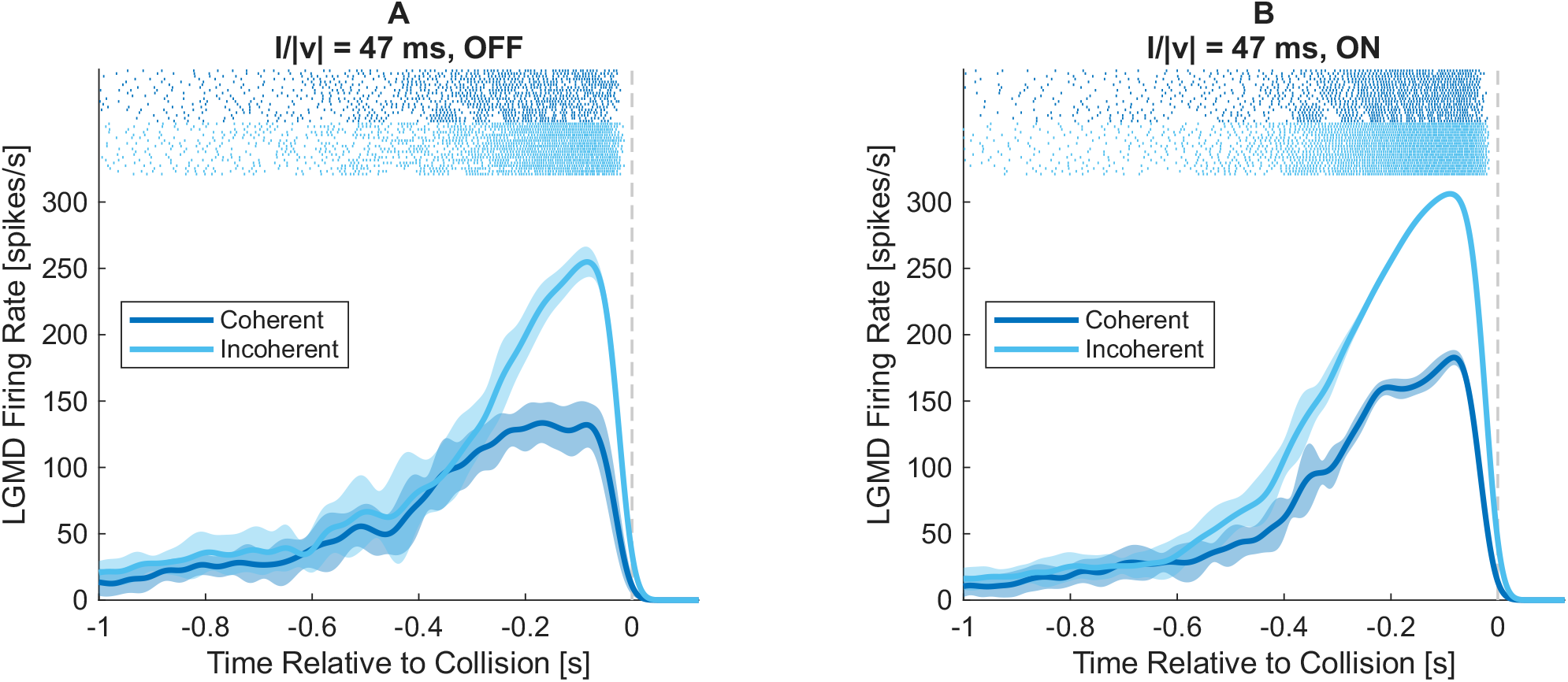
Comparison of modelled LGMD responses between coherent and incoherent looming stimuli. Model firing rates were obtained from a spike histogram with 1 ms bin width, smoothed by a Gaussian convolution with standard deviation of 21 ms. Lines and shaded regions are the mean and standard deviation of firing rates across model runs (*n* = 19), respectively. Dashed vertical gray lines indicate the time of collision. Spike rasters for each individual trial are given at the top of each plot. **A** shows a comparison of mean LGMD firing rate between coherent and incoherent OFF looming stimuli with *l/* | *v*| = 47 ms, while **B** shows the comparison of equivalent coherent and incoherent ON looms.

For ON stimuli, experimental data indicate that the effect of coherence on LGMD firing rate should be minimal [16]. However, the model once again exhibits a preference for incoherent stimuli (Fig 7B). This preference becomes apparent earlier in the loom than for OFF stimuli, and results in an overall larger increase in firing—a 58% increase in total spike count, on average.

While the preference of the model for incoherent stimuli is discouraging, it is interesting to note that this effect is lessened for OFF stimuli. This would suggest that some mechanism of coherence selectivity has a greater effect during OFF stimuli than ON, which would correspond to in vivo observations—even if said mechanism is not sufficiently powerful to create true coherence selectivity for OFF stimuli in the model.

To investigate this point, it is necessary to separate what is happening in the LGMD from what is happening in the rest of the model: the lamina, medulla, and lobula. This can be done by examining the behaviour of the direct excitatory inputs to the LGMD, which are the cumulative result of all prior elements of the network. Fig 9 shows the summed firing of the two sources of excitation to the LGMD: the TmAs (supplying input to field A), and the excitatory DUB neurons (supplying input to field C). For both of these neuron types, a preference for incoherent stimuli is readily apparent over the entire course of the looming stimulus, and is far greater in magnitude than for the LGMD. For OFF stimuli, the TmAs exhibited a 210% increase in spiking for incoherent stimuli over coherent, while ON stimuli showed a 209% increase. The excitatory DUB neurons exhibited a similar increase in spiking of 187% for incoherent ON stimuli relative to coherent ones (OFF stimuli generated negligible DUB response and were omitted from analysis). This could suggest that some mechanism is in the model is partially shielding the LGMD from the strong *incoherence* selectivity of its inputs, and that the incoherence selectivity seen in the modelled LGMD is a result of factors in its afferents, rather than the LGMD itself.

**Fig 9.**
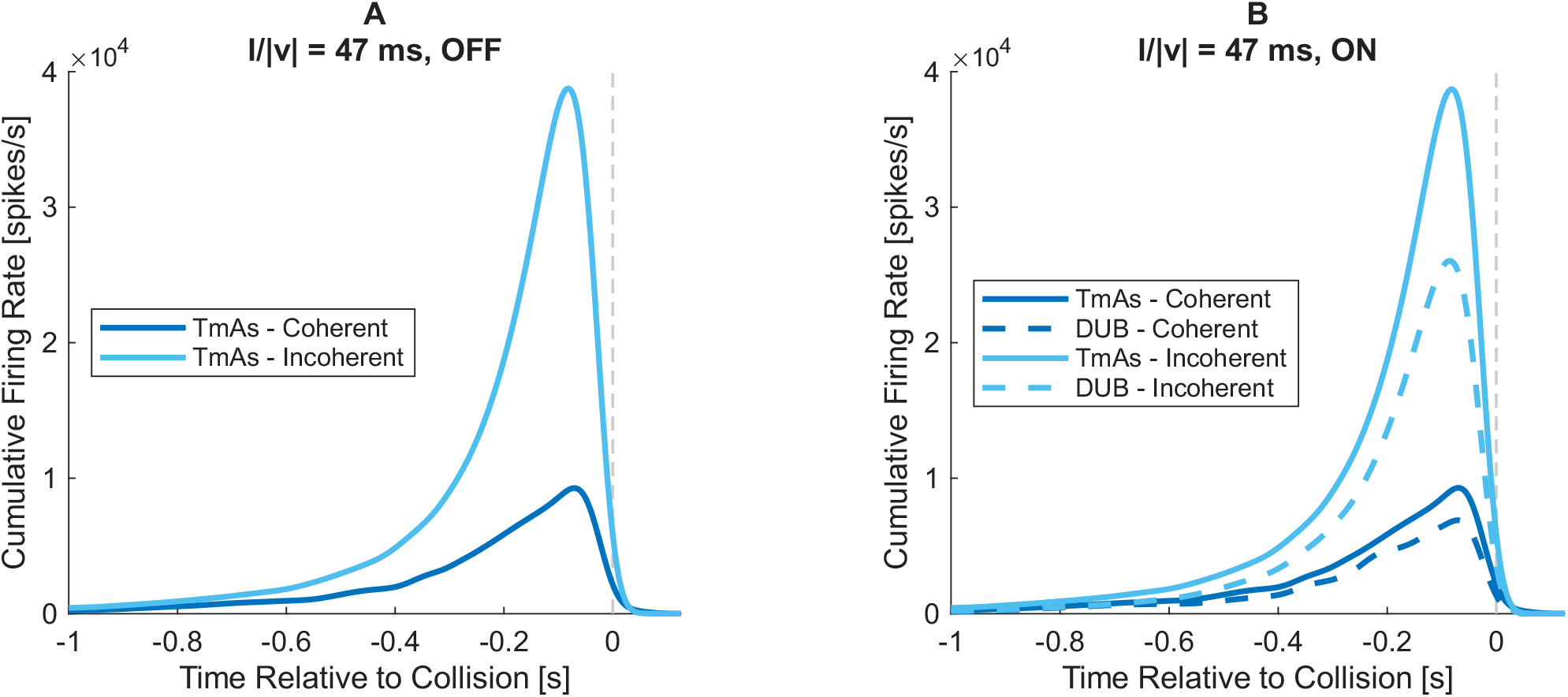
Comparison of excitatory inputs to the LGMD between coherent and incoherent looming stimuli. Model firing rates were obtained from a spike histogram with 1 ms bin width, smoothed by a Gaussian convolution with standard deviation of 21 ms. Dashed vertical gray lines indicate the time of collision. Firing rates are the cumulative sum of all neurons of a given type, and are averaged across multiple model runs (*n* = 19). **A** shows a comparison of cumulative TmA firing rate between coherent OFF looms with *l/*|*v*| = 47 ms and their incoherent equivalents. **B** compares the firing rates of both TmAs and excitatory DUB neurons across coherent and incoherent ON looms (*l/*|*v*| = 47 ms).

While greater activity in excitatory afferents seen during incoherent stimuli would naturally produce a greater response from the LGMD, the model assigns these neurons additional roles beyond providing the LGMD with input. Namely, the TmAs serve as the input to the OFF and ON FFI neurons, based on the modelling work from Olson et al. [31] and anatomical data from Wernitznig et al. [37]. Thus, increased activity of the TmAs and excitatory DUBs should be accompanied—and perhaps mitigated—by increased feedforward inhibition. Fig 10 shows this increased activation of feedforward inhibition during incoherent stimuli for both OFF and ON cases, with OFF FFI neurons showing a 192% increase in spiking in response to their corresponding stimuli, and ON FFI neurons showing a 166% increase. These concurrent increases in both excitation and inhibition may create the partial shielding of the modelled LGMD from the incoherence selectivity of its inputs. Also note that while ON and OFF TmAs showed a nearly identical increase in spiking as stimuli became incoherent, the increase in feedforward inhibition during OFF stimuli was prominently larger. This may be part of why OFF stimuli generated less incoherence selectivity in LGMD spiking.

**Fig 10.**
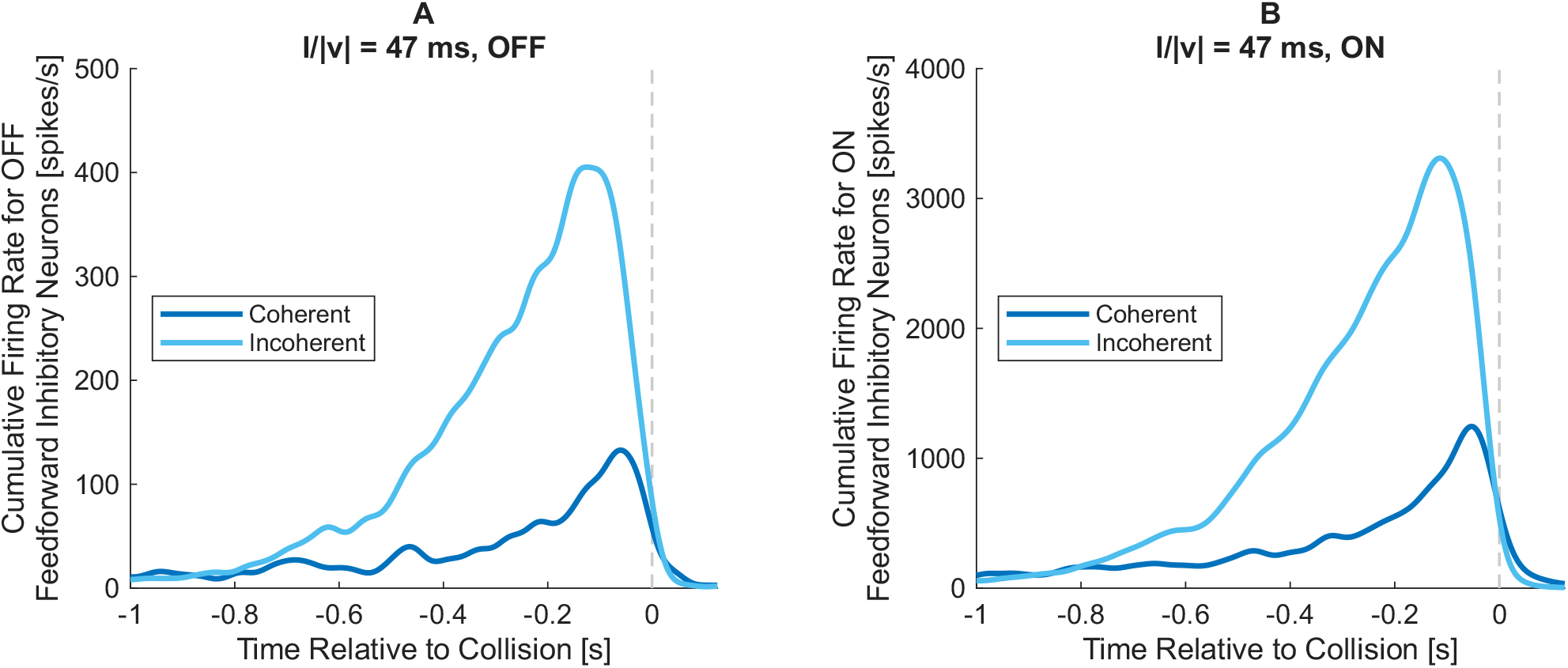
Comparison of feedforward inhibition to the LGMD between coherent and incoherent looming stimuli. Model firing rates were obtained from a spike histogram with 1 ms bin width, smoothed by a Gaussian convolution with standard deviation of 21 ms. Dashed vertical gray lines indicate the time of collision. Firing rates are the cumulative sum of all neurons of a given type (either OFF or ON FFI neurons, depending on the stimulus), and are averaged across multiple model runs (*n* = 19). **A** shows the response of OFF FFI neurons during coherent OFF looms with *l/* | *v*| = 47 ms, as well as during equivalent incoherent stimuli. **B** shows the same comparison for ON FFI neurons during ON stimuli.

Another factor to be considered is the relative timing of excitation and inhibition to the LGMD. For incoherent stimuli, all of these became earlier. The peaks of mean TmA firing rates during OFF and ON stimuli occurred 11 and 12 ms earlier, respectively, for incoherent stimuli, while the peak of the mean firing rate for excitatory DUB neurons during ON stimuli occurred 14 ms earlier. OFF and ON FFI neurons, however, had the peaks of their mean responses shifted 65 and 60 ms earlier when their respective stimuli were made incoherent. This relatively earlier peaking of inhibition during incoherent stimuli may partially explain why increased sensitivity to incoherent stimuli becomes most visible late in the loom, especially for the OFF stimulus (see Fig 8).

Another factor which should compensate for incoherence selectivity in excitation to the LGMD are its active conductances. The HCN and K_D-like_ conductances in field A contribute to the actual LGMD neuron exhibiting a preference for coherent stimuli [32]. To examine this in the modelled neuron, the method for generating incoherent stimuli was changed. Instead of mapping ommatidial luminance values to random sites (per Fig 5), the “axons” of the modelled TmAs were remapped in a similar fashion. This was done by shuffling TmA synaptic outputs to random sites, prior to simulating their branching to adjacent sites on field A. As TmA inputs to non-LGMD neurons (i.e. FFI and excitatory DUB neurons) were not remapped, this focuses the question of coherence selectivity solely on factors within the LGMD. The resulting LGMD looming responses are shown in Fig 11.

**Fig 11.**
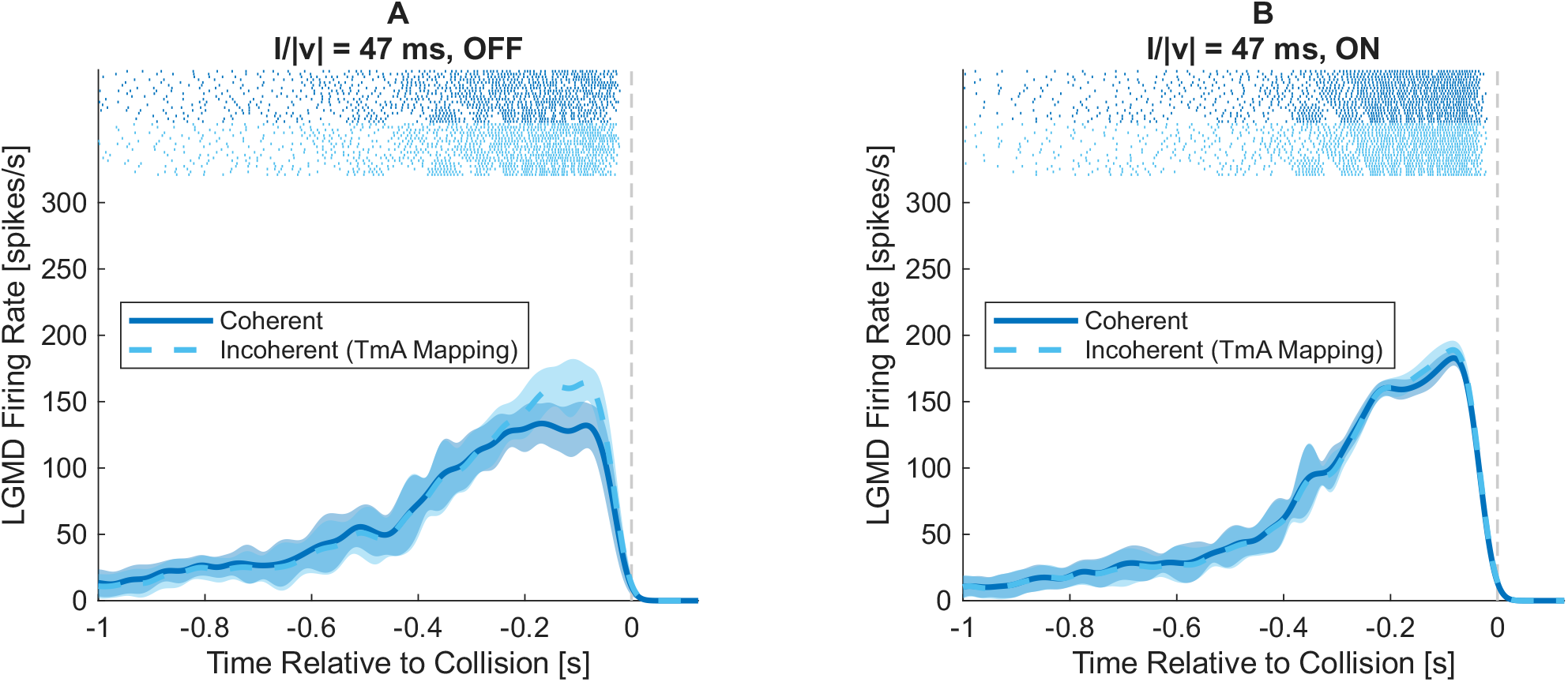
Responses of the LGMD model to coherent and incoherent mapping of its field A inputs. Model firing rates were obtained from a spike histogram with 1 ms bin width, smoothed by a Gaussian convolution with standard deviation of 21 ms. Lines and shaded regions are the mean and standard deviation of firing rates across model runs (*n* = 19), respectively. Dashed vertical gray lines indicate the time of collision. Spike rasters for each individual trial are given at the top of each plot. **A** shows a comparison of mean LGMD firing rate with both coherent and incoherent TmA mappings, during OFF looming stimuli with *l/*|*v*| = 47 ms. **B** shows a similar comparison for equivalent ON looms.

In these results, LGMD firing rate appears mostly insensitive to TmA mapping coherence, especially for ON stimuli. For OFF stimuli, the LGMD continues to exhibit a slight preference for incoherent stimuli over a window starting ∼250 ms before collision, up to ∼50 ms. Prior to this window, mean firing rates are equal, or very slightly reduced for incoherent stimuli. As shown via modelling by Dewell and Gabbiani [32], this result is nontrivial; a purely passive field A would actually exhibit a preference for incoherent stimuli, due to coherent inputs driving local membrane potentials closer to the reversal potential of excitatory synapses and reducing input currents. Thus, the lack of a coherence preference over most of the duration of the looming stimuli likely indicates that the modelled passive conductances have some effect in shifting the preference of the LGMD toward coherent stimuli—though not sufficiently to produce notable coherence selectivity.

Based on Fig 9 and Fig 11, it can be said that the incoherence selectivity of the complete model stems primarily from elements presynaptic to the LGMD itself. This raises the question: what element is responsible? Given that lateral excitation (which was omitted from the model) is important to the coherence selectivity of LGMD response in vivo [33], the inverse assumption might be made that lateral inhibition conveys a preference for incoherent stimuli. This is logically sound, as adjacent stimuli forming a coherent edge would be capable of mutual lateral inhibition, while randomly-scattered elements of an incoherent stimulus are far less likely to fall within the limited range [30] of each others’ generated inhibition. To test this assumption, basic coherent and incoherent looming responses were reexamined with lateral inhibition removed, as presented in Fig 12.

**Fig 12.**
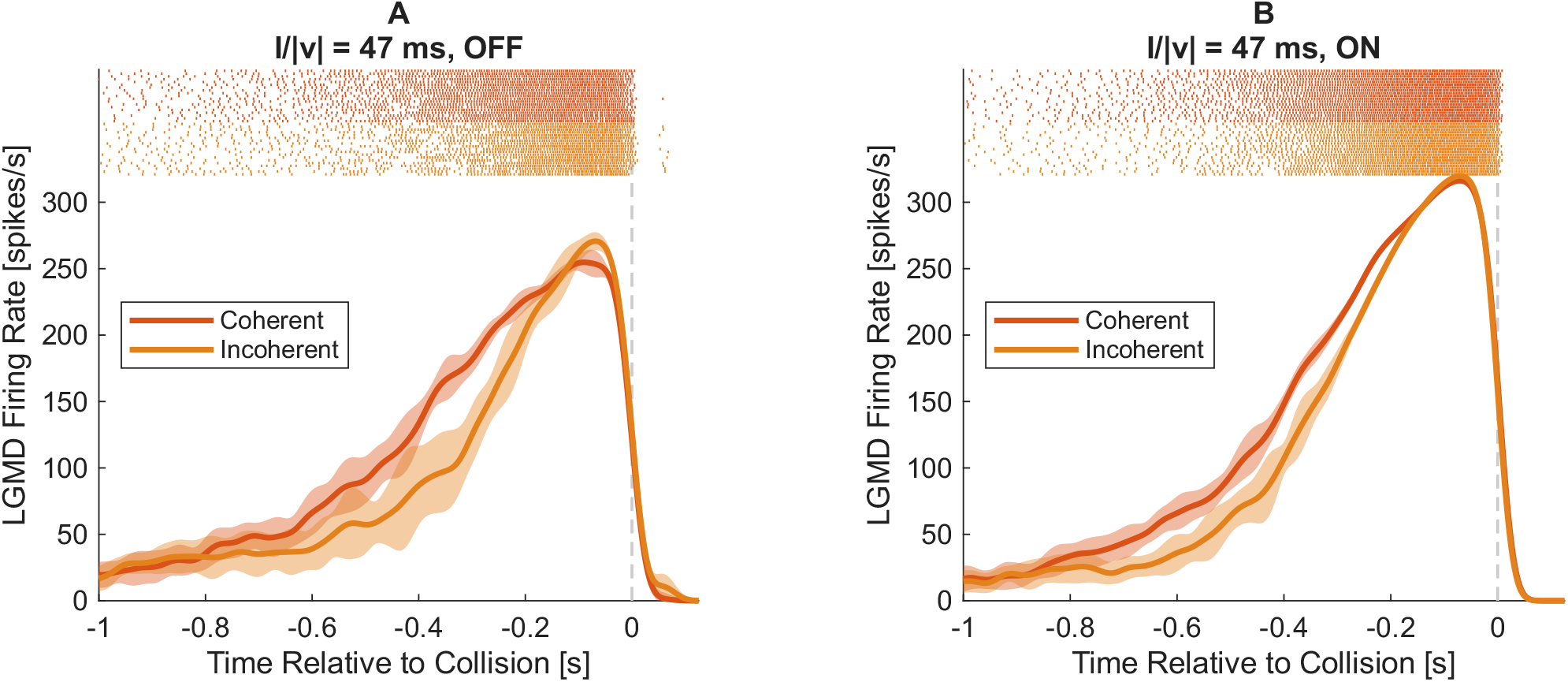
Comparison of modelled LGMD responses between coherent and incoherent looming stimuli, after removing lateral inhibition. Model firing rates were obtained from a spike histogram with 1 ms bin width, smoothed by a Gaussian convolution with standard deviation of 21 ms. Lines and shaded regions are the mean and standard deviation of firing rates across model runs (*n* = 19), respectively. Dashed vertical gray lines indicate the time of collision. Spike rasters for each individual trial are given at the top of each plot. Lateral inhibitory connections were removed, per Fig 5. **A** shows a comparison of mean LGMD firing rate between coherent and incoherent OFF looming stimuli with *l/* | *v*| = 47 ms, while **B** shows the comparison of equivalent coherent and incoherent ON looms.

Contrary to prior results, the LGMD here exhibits an overall preference for coherent stimuli, with incoherent stimuli showing 13% reductions in spike count for both OFF and ON stimuli. This coherence preference disappears around the peak time for LGMD firing rate. The reasons for this are likely the same factors which create a stronger incoherence preference around this time in the prior cases examined.

While removing lateral inhibition explains the disappearance of incoherence preference, it does not explain the emergence of a coherence preference, as the modelled LGMD exhibits negligible coherence preference of its own (per Fig 11). Removing lateral inhibition removes the only element in the input network up to the TmAs which exhibits any dependence on the spatial location of individual ommatidial inputs. This leaves only wide-field medulla neurons, such as those governing feedforward inhibition, as being capable of involvement. The behaviour of these neurons—namely, the OFF and ON FFI neurons—is shown in Fig 13.

**Fig 13.**
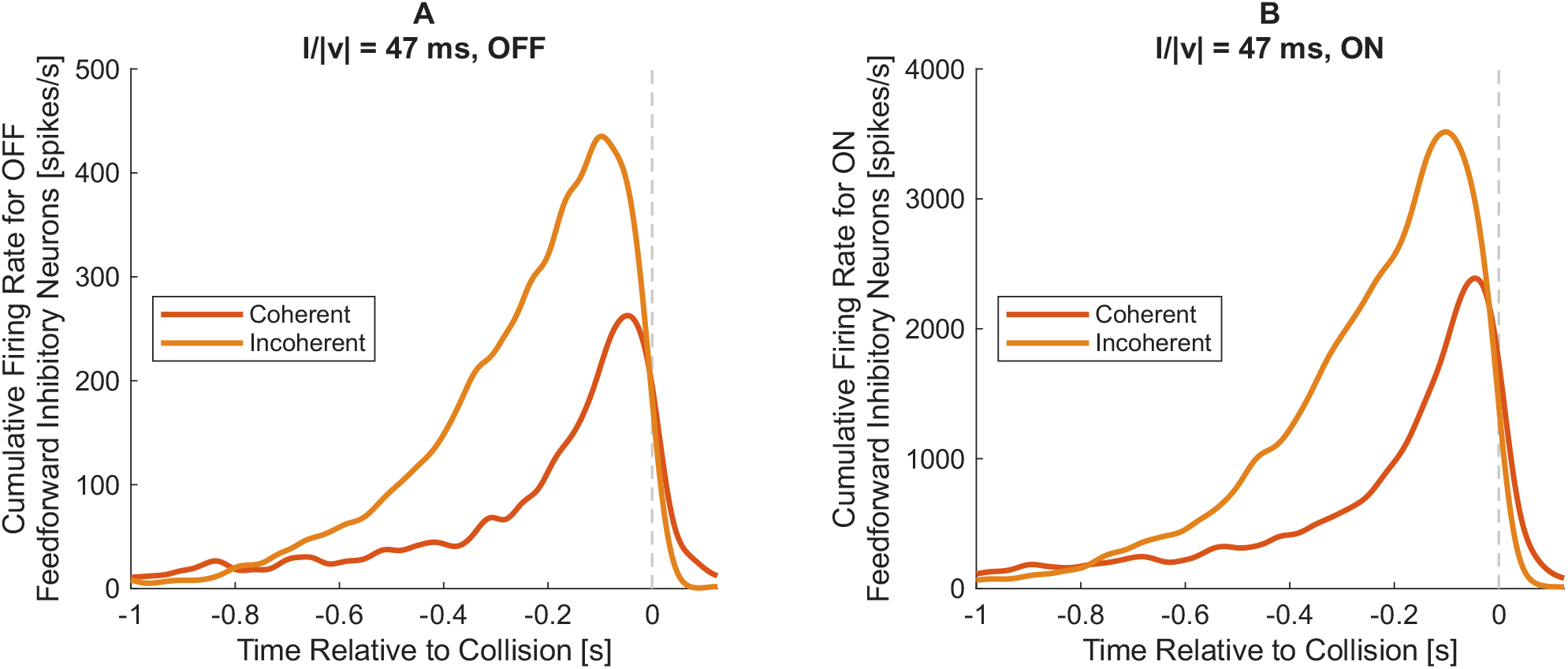
A comparison of feedforward inhibition to the LGMD between coherent and incoherent looming stimuli, after removing lateral inhibition. Model firing rates were obtained from a spike histogram with 1 ms bin width, smoothed by a Gaussian convolution with standard deviation of 21 ms. Dashed vertical gray lines indicate the time of collision. Firing rates are the cumulative sum of all neurons of a given type (either OFF or ON FFI neurons, depending on the stimulus), and are averaged across multiple model runs (*n* = 19). Lateral inhibitory connections were removed, per Fig 5. **A** shows the response of OFF FFI neurons during coherent OFF looms with *l/* | *v*| = 47 ms, as well as during equivalent incoherent stimuli. **B** shows the same comparison for ON FFI neurons during ON stimuli.

Surprisingly, the FFI neurons still exhibit a preference for incoherence, even when lateral inhibition is removed from their inputs. OFF FFI neurons generate 84% more spikes when their preferred (OFF) looms are made incoherent, while ON FFI stimuli exhibit a 70% increase in spike generation under similar conditions for simulated ON looms. These are lesser increases than when lateral inhibition was present, but are still prominent ones. This near-doubling of direct inhibition to the LGMD during incoherent stimuli likely accounts for the majority of the coherence selectivity observed in Fig 12. The increased feedforward inhibitory activity still exhibits a shift in its peak time after removing lateral inhibition, with the peak of mean firing rate occurring 51 ms earlier for OFF FFI neurons during OFF looms, and 56 ms earlier for ON FFI neurons during ON looms. This may be a factor in why coherence selectivity seems time dependent over the course of the loom, although the further tests would be required to explore this fully.

### Complex backgrounds and the roles of inhibition types

Stimuli incorporating wide-field background motion provide another opportunity to reference the model against experimentally-observed responses. They also provide an opportunity to identify roles of different types of inhibition in shaping the response of the LGMD—roles which are often best elucidated by removing various inhibition types.

Fig 14A shows the LGMD response to an OFF looming stimulus against a blank background, both for the complete model and with the various inhibitory mechanisms removed. While removing global, lateral, and feedforward inhibition all increase the LGMD firing rate, the size and temporal distribution of these increases differ. Removing global inhibition generates the largest increase in peak firing rate, with the mean LGMD firing rate across trials reaching 383 spikes/s (compared with 136 spikes/s for the baseline model). This peak occurs near the the time of collision, notably later than in the baseline model, suggesting that global inhibition is at its most important in the latter part of a looming stimulus. However, this might not be entirely true. Given that global inhibition in this model occurs prior to the divergence of feedforward inhibitory pathways, as suggested by the work of Olson et al. [31], its removal increases not only LGMD excitatory input, but also feedforward inhibitory activity. Thus, the particular change in firing rate trajectory seen after removing global inhibition may not be solely attributable to the properties of global inhibition alone, but also to how its removal differentially affects the behaviour of the LGMD and the FFI neurons. This requires further examination to elucidate, but should be borne in mind when examining effects of the removal of global inhibition, along with lateral inhibition (which is also included in inputs shared between the LGMD and FFI neurons here).

**Fig 14.**
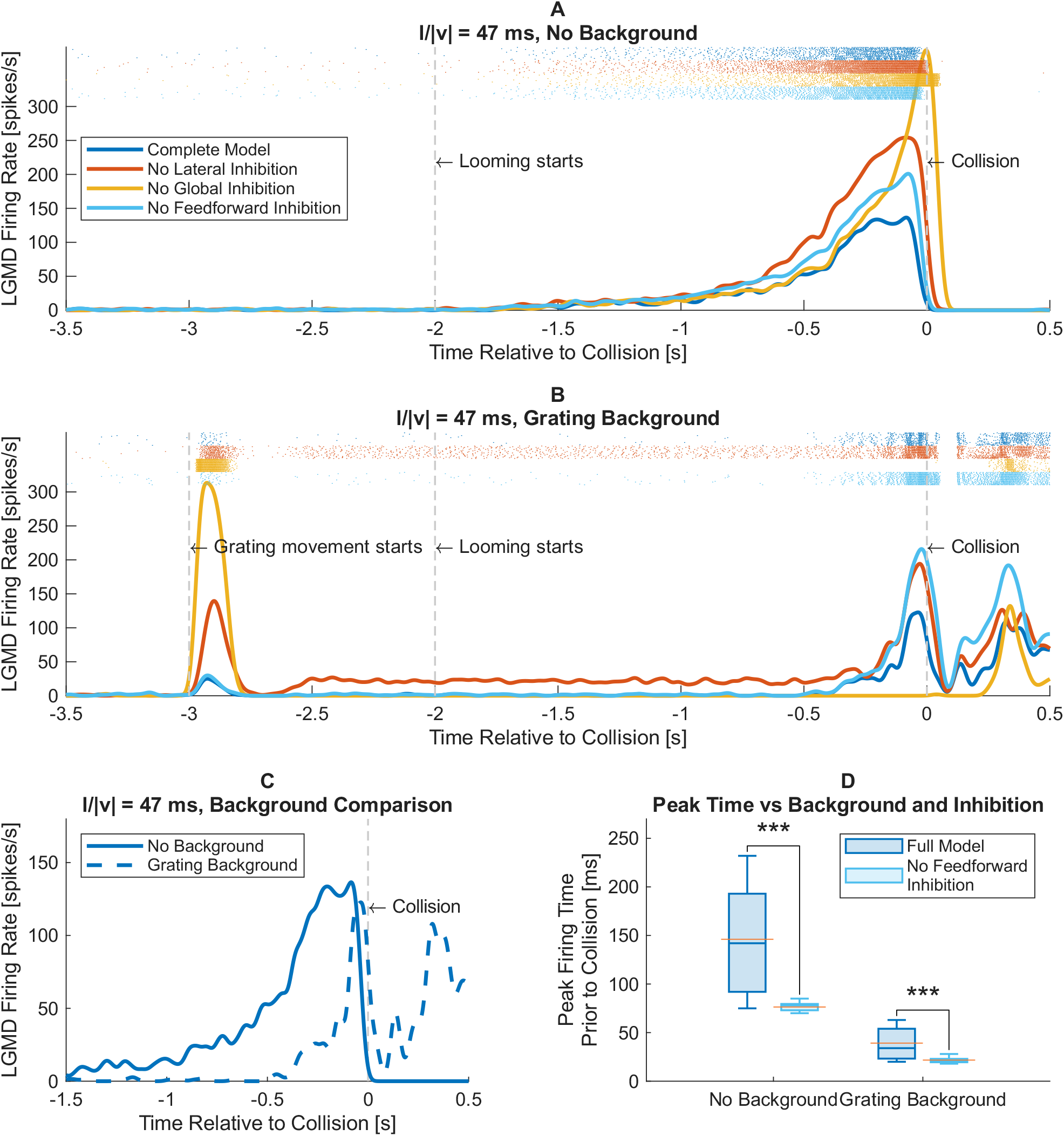
Effects of different backgrounds and removal of inhibitory mechanisms on modelled LGMD responses. Model firing rates were obtained from a spike histogram with 1 ms bin width, smoothed by a Gaussian convolution with standard deviation of 21 ms. Firing rates are averaged across multiple model runs (*n* = 19). Dashed vertical gray lines indicate start times for grating motion and looming, as well as time of collision. Spike rasters for each individual trial are given at the top of each plot. **A** shows the LGMD firing rate of various model configurations to basic OFF looms with *l/* | *v*| = 47 ms. Configurations include the complete model, as well as versions without global, lateral or feedforward inhibition. **B** shows responses of the same model configurations to an identical looming stimulus against the background of a moving sinusoidal grating. **C** directly compares the LGMD firing rate of the intact model between looming stimuli with and without the grating background. **D** shows the effect that removal of feedforward inhibition has on the time of peak firing rate relative to collision via a box plot. In addition to indicating the median peak time for each condition, the mean values (orange lines) are also given for reference. A Mann-Whitney U test showed that removal of feedforward inhibition had a significant effect on the time of LGMD peak firing rate during looming, both with no background present (*U* (19, 19) = 19, *p <* 0.001) and in the presence of a moving grating background (*U* (19, 19) = 41, *p <* 0.001).

Removals of both lateral and feedforward inhibition create less distinctive shifts in the temporal distribution of LGMD spiking, with the underlying shape largely preserved. The removal of feedforward inhibition in particular does appear to result in the creation of a sharper peak in the mean LGMD firing nearer to the time of collision, but further exploration, given below, is needed to make their roles clearer.

Fig 14B shows the responses of the same model configurations to an identical looming stimulus accompanied by a full-field sinusoidal grating background which begins moving before the loom starts. These full-field motions do not trigger sustained responses from the LGMD [30], but do affect its responses to looming [27]. The intact model exhibits minimal LGMD response to the beginning of grating motion, reaching a peak of only 25 spikes/s. It also maintains a response to looming, albeit with changes to the firing rate trajectory. Fig 14C directly compares the looming response of the model with and without a moving background. The average peak time shifts closer to collision when the background is present, from 146 ms before collision to 39 ms before collision. The rise phase also becomes substantially later. Both of these effects are qualitatively consistent with observations from Silva et al. [27]; however, as they used different parameters for both looms and gratings, direct quantitative comparisons cannot be made.

Another apparent change to the LGMD response in the presence of a grating is substantial post-collision firing. This is apparent in Fig 14B for not only the full model, but for all tested inhibitory configurations as well. This is likely an artefact of the looming stimulus used. As the looming stimulus stops moving on the last timestep before collision, its final subtense angle is 179°, leaving a small sliver of the grating background exposed near the edge of the eye. This transforms the small visible portion of each grating stripe into a translating small-field stimulus, which presynaptic lateral inhibition does not fully suppress in vivo [30]. Unlike the biological LGMD [30], the model exhibits no habituation to such stimuli, and presumably would continue firing if left in this state.

Returning to Fig 14B, it can be seen that removing lateral inhibition causes a sustained response to wide-field motion, which is not seen for any other model structure tested. However, it is not the only inhibition type whose removal has a different effect when a moving background is present. Unlike a standard loom, for which the removal of global inhibition creates much higher LGMD firing rates, the removal of global inhibition when wide field motion is present abolishes the model’s looming response entirely, even though the response to initiating grating motion is substantially increased.

While lateral inhibition has a clear role in suppressing responses to wide-field background motion, and global inhibition likely has multiple roles, the role of feedforward inhibition is less clear at first glance. To illuminate it further, the time of peak firing rate for both stimulus types (blank and grating backgrounds) was examined both with and without the inclusion of feedforward inhibition, shown in 14D. As seen here, feedforward inhibition has a highly significant (*p <* 0.001) and relatively consistent effect on the peak time of LGMD firing rate. Removing feedforward inhibition reduces the mean time between peak firing rate and collision to 52% of its baseline value for stimuli with no background, and 55% of its baseline value for stimuli with a moving grating background.

Beyond changing when peak firing occurs, the removal of feedforward inhibition substantially reduces its variability. After removing feedforward inhibition, the standard deviation in peak time reduces to 8% of its former value for stimuli with no background, and 18% for stimuli with a grating background. Thus feedforward inhibition, as modelled, increases the variability of peak time in addition to shaping its location.

### Diverse object trajectories

While the LGMD responds preferentially to looming, it also responds to other motion by small objects, such as translation [23, 27, 30]. Translating stimuli provide opportunities to examine modelled LGMD responses across varying parts of the visual field, and to confirm the model’s preference for looming. Moreover, characteristic responses after transitions from translation to looming or vice-versa provide points of reference for the model’s behaviour.

Fig 15 shows responses to a variety of stimulus trajectories either taken directly from work by McMillan and Gray [23], or chosen to have equivalent velocities and location for trajectory change (for the looming-to-translation case). Much like experimental data, the model rapidly transitions to a firing rate corresponding to the new trajectory of object motion after directions changes. When shifting from looming to translation, however, there is a brief peak in firing rate not visible in either of the equivalent translating or looming stimuli. This phenomenon is consistent with experimental observations for transitions from looming to translation using other trajectory geometries [23]. However, the transition from translation to looming does not exhibit a corresponding temporary drop in firing rate, instead transitioning smoothly from translation to looming firing rates. The comparison to data from McMillan and Gray [23] in Fig 15B3 shows this clearly; in fact, the model LGMD firing rate appears to rise sharply just before the drop seen in experimental data would occur. This rise in firing rate also begins prior to the translation to looming itself, and is not clearly visible in the corresponding looming response (Fig 15B2).

**Fig 15.**
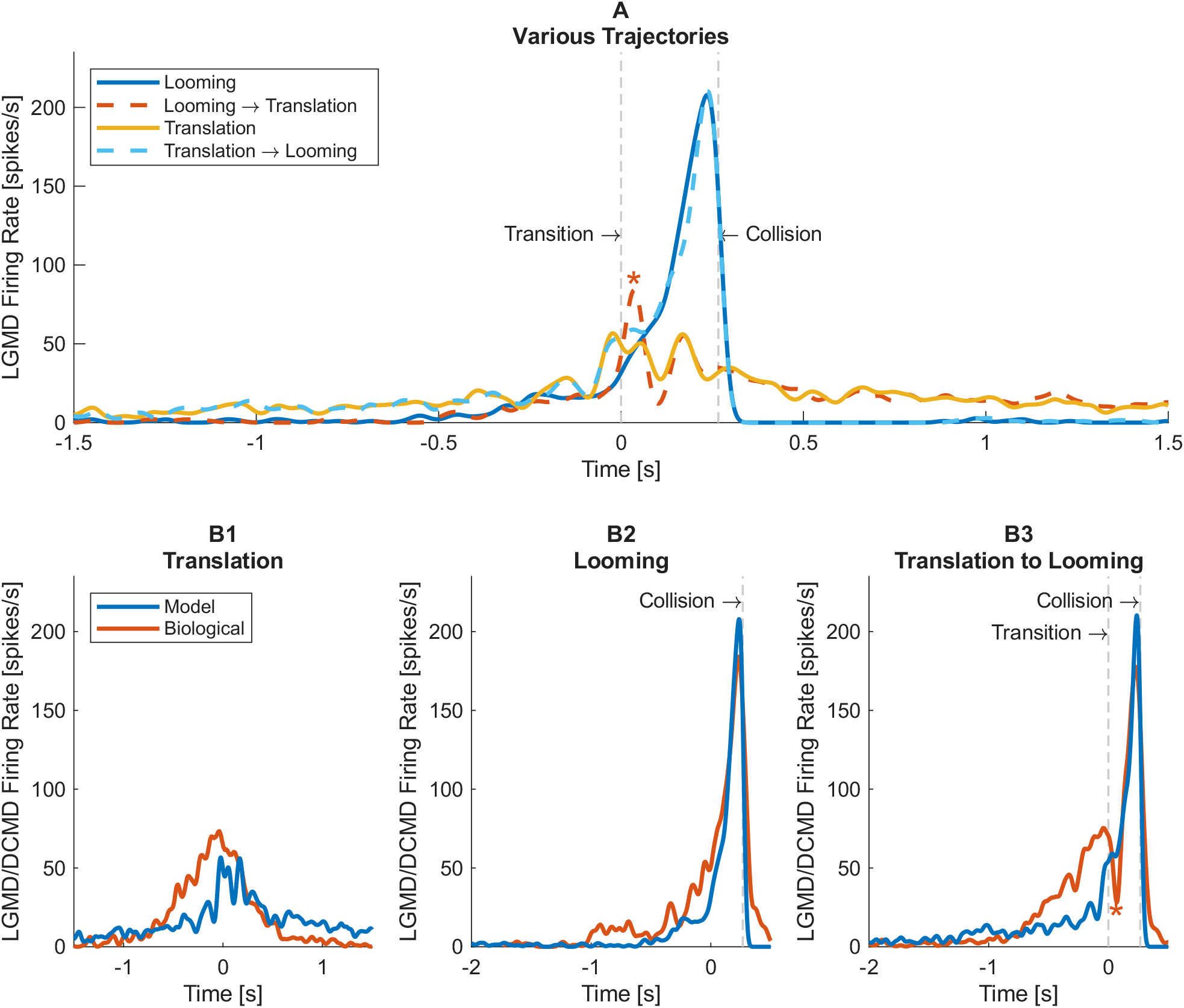
Responses of the modelled LGMD to stimuli of varying trajectories, compared with data from McMillan and Gray [23]. Model firing rates were obtained from a spike histogram with 1 ms bin width, smoothed by a Gaussian convolution with standard deviation of 21 ms, for comparability to the 50 ms FWHM Gaussian used in the biological data. Model firing rate data were averaged across multiple model runs (*n* = 19), while experimental values were averaged across multiple animals (*n* = 20). All time values were aligned to the time (*t* = 0) at which the stimulus reached a point 80 cm away along the central axis of the eye. Vertical dashed grey lines indicate transitions in stimulus trajectory, along with collisions. The stimulus consisted of a 3.5 cm radius disc moving at a speed of 3 m/s, equating to *l/* | *v*| = 12 ms during looming portions. **A** compares modelled LGMD firing rate for looming, translating, translating-to-looming and looming-to-translating stimuli. A local peak in response to transitioning from looming to translation, typical of biological responses to similar stimuli [23], is indicated via an asterisk (*). **B1-3** show comparisons of modelled LGMD firing rate responses to translating (**B1**), looming (**B2**), and translating-to-looming (**B3**) stimuli to the corresponding DCMD response data from McMillan and Gray [23]. In **B3**, the characteristic drop in firing rate observed experimentally after transitioning from translation to looming is indicated with an asterisk (*); note its absence in the model response.

The translation response of the model (Fig 15B1) exhibits a sharp increase in firing rate as the stimulus reaches the central axis of the eye (where the transition to looming would occur in Fig 15B3), in contrast with the more gradual increase seen in the experimental data. As such, that element of the translation-to-looming response derives from the translation portion. Interestingly, up until *t* ≈−0.7 s, the model slightly overestimates LGMD firing rate. It is only just prior to the object reaching the center axis of the eye that the model firing rate again approaches experimental data. It then follows experimental behaviour until around *t* ≈ 0.5 s, after which the model again overestimates firing rate.

While the the model has issues replicating translating responses and the transition from translation to looming, it exhibits a strong preference for looming stimuli, in terms of firing rate. As this is a key aspect of LGMD behaviour [25, 63], such a preference does reinforce that at least some aspects of its underlying principles are correct.

## Discussion

While the results show differences between the responses of the LGMD model and its biological counterpart, there are many similarities, including

- the general response to looming stimuli;
- the linear relationship between peak response time and the stimulus parameter *l/*|*v*|;
- the insensitivity of the LGMD to wide field motion;
- the ability of the LGMD to discriminate small-field motion such as looming against moving backgrounds, with the rise and peak time of the looming response exhibiting delays compared to responses with no background present;
- the transition of LGMD responses to an appropriate new firing rate trajectory after transitions in stimulus behaviour; and
- the preference of the LGMD for looming over translating motion.

The fact that the model replicates multiple aspects of LGMD behaviour well indicates the plausibility of assumptions underlying its construction. However, its discrepancies from its biological counterpart provide useful points of reference for future improvement of the model. They also highlight the importance of biological mechanisms specifically omitted from the model during its construction.

### Structure of LGMD inputs

Olson et al. [31] observed that a successful model of feedforward inhibitory neurons in the DUB incorporated specific neurons (the LMCs and TmAs) and general mechanisms (global and lateral inhibition) understood to be present in the inputs to the LGMD. This model demonstrates that not only do the FFI DUB neurons and LGMD likely both have similar computational mechanisms in their input networks, but that it is plausible for these neurons to share an input network up to and including the TmAs. This result is particularly significant when considering that the presynaptic network to the LGMD (for OFF looms—this excludes the excitatory DUB neurons) had its parameters fit solely through examination of the OFF FFI neuron responses, with the LGMD looming response being fitted solely through adjustment of the LGMD itself.

Moreover, producing realistic looming responses to ON stimuli only required adjustment of the excitatory DUB neurons. If these neurons did not share common inputs, one might expect that their different computational needs would have to be balanced through compromises when adjusting the properties of their shared input network during the parameter fitting process. The fact that the input network to three different types of modelled neuron could be adequately fit based on the responses of just one of them, while still allowing the other two to produce appropriate responses, further strengthens the proposition that these neurons share inputs in the biological case, though it remains to be tested further.

This overlap in inputs results in a structurally parsimonious network, with the potential to reduce energy costs to the animal. Based on recent observations regarding the nature of excitatory inputs during ON looms [16], such factors seem to be an evolutionary consideration elsewhere in the LGMD circuit. This means that there is a distinct evolutionary pressure toward the proposed network structure.

Beyond shared inputs for the LGMD and OFF FFI neurons, this model incorporates two other key structural assumptions:

- the existence of separate OFF and ON TmAs; and
- the dependence of excitatory DUB neurons on TmA input.

The ability of the resulting excitatory afferents to the LGMD to generate realistic OFF and ON looming responses shows that their structure is plausible, but does not provide confirmation of it. However, this result must also be taken in the context of the observations used to justify the model structure, namely:

- the existence of two different TmA morphologies [37, 41];
- input differences between these morphologies, including possible direct ON input to one but not the other [37];
- differences in the number of connections made at the LGMD [37]; and
- correspondences between the number of connections made by the two different TmA types and the relative strengths of ON and OFF field A input [16].

Taken together, the success of the model and the multiple observations from biological data suggest a reevaluation of TmA functionality is warranted.

One notable quirk of the model is the higher peak firing rate seen for an ON loom relative to an OFF loom at the same *l/* |*v*|, which is inconsistent with biological data [12, 16]. This difference comes not from the model overestimating peak firing rate for ON looms, but rather underestimating it for OFF looms. Part of this underestimation likely stems from the fitting process, coupled with the broadness of the peak in mean firing rate for OFF stimuli. This broadness traces back to the variability in peak times during OFF looms, which is greater than that observed biologically (Fig 7). The variability of peak time observed for modelled ON looms is lower than either model or biological data for OFF looms, with a standard deviation of 11 ms in modelled ON looms, compared to 54 ms for modelled OFF looms and 27 ms for data from Stott et al. [1].

The fact that biological peak time variability for OFF looms lies between the model variability for OFF looms and ON looms may indicate part of the reason why the ON looms show less peak time variability, and also reflect an omitted factor in the model. During an ON loom, much of the excitation to the both the model and the biological LGMD [16] arises from excitatory DUB neurons giving input to field C. In the model, the presence of this excitation during ON looms only may partially explain their less variable peak time. This idea is supported by the input to field C having less electrotonic separation from the SIZ, and thus a proportionally stronger influence on spike generation (see modelling by Dewell et al. [16]).

In this model, the excitatory DUB neurons projecting to field C are purely ON units; however, experimental data from Dewell et al. [16] showed that they provide some—albeit lesser—excitation to field C during OFF stimuli as well. The omission of field C OFF excitation would explain both why the model’s peak time for OFF responses is more variable than in biological data, and could also explain why it underestimates peak firing rate at lower *l/* |*v*| values.

Adding OFF excitation to field C could be implemented in multiple ways, structurally. One would be to incorporate excitation from the OFF TmAs into the inputs of the excitatory DUB neurons. Both experimentally observed TmA types have output synapses in the proximal medulla [37], and the neurons of the DUB form dendritic arbours there [14], making this idea biologically plausible. A second possibility is that the two TmA types are not purely ON and OFF cells, but are only predominantly of one polarity or the other. This would explain early observations suggesting that TmAs are ON/OFF cells [40].

One question not examined here is how the excitation to field C shapes the looming responses seen both in the model and in vivo. The characteristic response of the LGMD to OFF looms is understood to be dependent on the difference in stimulus properties encoded between its excitatory and inhibitory afferents, and the way they are shaped differently by field A and field C respectively [2, 15, 18]. Specifically, field A is understood to logarithmically compress excitatory inputs in a way that field C does not for inhibitory inputs. However, field C also receives excitation during looms, raising the question: what mechanisms ensure that field C excitation during ON looms is correctly shaped relative to feedforward inhibition?

One potential way of shaping DUB excitation to field C is through the behaviour of the neurons that generate it. Shared input between the LGMD, OFF FFI neurons and excitatory DUB neurons would place increased weight on how each neuron type processes this input to achieve different computational ends. The LGMD notably has an electrotonically extensive structure [13, 18] and prominent active conductances [32, 38] to accomplish this task, while Olson et al. [31] demonstrated how strong post-firing conductances can contribute to the OFF FFI neurons coding linearly for stimulus subtense angle, as they do in vivo [15]. While the firing of both excitatory DUB neurons and their ON TmA inputs is shown in Fig 9, no rigorous examination is given.

Another possibility is that the processing of excitatory and inhibitory inputs to field C is differentiated by their relative positioning within field C. As mentioned previously (see General LGMD structure and properties), inhibitory inputs to field C were moved to its proximal portion, which then had its axial resistance *R*_*a*_ reduced to further reduce its electrotonic distance from the SIZ. The excitatory inputs remained in the distal portion, which had a much higher *R*_*a*_ value, thus placing them at a greater electrotonic distance from the SIZ. Calcium fluorescence data show the location of DUB excitation in field C [16], but no comparison with the mapping of field C inhibition appears to exist—athough the lack of such investigation is understandable, as the discovery of separate excitatory and inhibitory neurons in the DUB is a recent one [16].

### Roles of inhibition types

The success of the model in replicating a variety of LGMD responses makes it an effective platform for examining the roles of different inhibition types in shaping these responses.

The role of lateral inhibition in the model is the simplest. As removing lateral inhibition is the only modification to the model which causes it to become sensitive to wide-field motion, it can be concluded that lateral inhibition is necessary to suppress the response to wide-field motion. This agrees with the role assigned to it by Olson et al. [31], as well as its originally hypothesized function [30]. In this model, the LGMD still exhibits a response to the looming stimulus. In vivo, however, an LGMD neuron would experience habituation of its excitatory inputs from this stimulus [30], which would sharply reduce its response to looming [64, 65]. Unlike habituation to small-field stimuli, this habituation would occur over the entire visual field, rendering the eye partially blind to looming stimuli. As such, the role of lateral inhibition in preventing wide-field motion from exciting the LGMD is more important than implied by the model.

The role of global inhibition is more nuanced. As its removal causes the highest peak firing rates for looming without a background, and this peak firing rate is positioned late in the loom, it can be inferred that it improves the dynamic range of the LGMD, allowing it to respond earlier in object approach without excess firing in the later stages. This is in agreement with the role proposed by Zhu et al. [33], and with what was shown in the looming responses of modelled DUB neurons by Olson et al. [31].

Removing global inhibition also causes the largest increase in the response to initiation of wide-field motion. This is a more prominent effect on the response to initiating grating motion than observed in modelling by Olson et al. [31], though the differences in the size of the grating area—40° across versus the 180° used here—may explain the difference, as may the difference between feedforward and feedback schemes for global inhibition.

It is in looming against a moving grating that the most unusual effect of global inhibition appears. While removal of global inhibition causes the most prominent *increase* in firing rate during looming against a blank white background, its removal during the presence of a grating causes the looming response to *disappear* entirely. This seemingly contradictory result can be explained by the relative positions and structures of different mechanisms within the network. Global inhibition occurs prior to lateral in the model, affecting both the signals that generate lateral inhibition and the signals which lateral inhibition acts upon. This means that global inhibition simultaneously

- reduces the amount of lateral inhibition required to suppress wide-field motion; and
- reduces the amount of lateral inhibition generated during wide-field motion.

Thus, when global inhibition is removed, the resultant increase in feedforward lateral inhibition suppresses all excitatory input to the LGMD, including that generated by looming. Reducing the requisite lateral inhibition is especially important given that it is both delayed and persistent, a factor found to be required for it to create biologically plausible responses in work by Olson et al. [31]. As such, lateral inhibition sufficient to block inputs from grating motion at their full strength would also persist into looming and block response to it entirely as well—exactly as seen here when global inhibition is removed.

Another important element of global inhibition is that it is *global*, and thus governed more strongly by wide-field motion than small-field stimuli, even if the small-field stimuli generate stronger inputs at the level of individual ommatidia (i.e. larger or faster changes in local contrast). This means that once the small-field stimuli produce sufficiently strong local excitation, global inhibition should permit them to stand out further against background motion. The loss of this effect upon removing global inhibition, coupled with the aforementioned increase in lateral inhibition, results in any response to looming being completely lost.

It should be noted that the function of global inhibition in enhancing looming response when a moving background is present appears to be enhanced by its feedback configuration. When using the feedforward-style global inhibition of Olson et al. [31] in developing this model, it was found that tunings of it which were sufficient to normalize looming responses against a blank white background would also invariably abolish them once a moving background was introduced. Conversely, a feedback inhibitory system is inherently self-limiting, and the resulting inhibition cannot completely abolish the response of the system, regardless of stimulus, meaning that some response will always be preserved. Note that this feedback configuration was enabled by the addition of the IMUs between the LMCs and TmAs; this provides one potential role for the neurons presynaptic to the TmAs identified by Wernitznig et al. [37]. This particularly relevant for TmA2, for which some of these neurons likely represent intermediaries between the LMCs and the TmA.

Contrary to global inhibition, the included lateral inhibition must be feedforward to perform its function of completely silencing responses to wide-field motion, rather than merely scaling them. This makes the order of the two mechanisms of particular importance; the functions of global inhibition become irrelevant if persistent lateral inhibition from wide-field motion has already silenced excitation upstream. The difference in requisite structure for global and lateral inhibition to perform their functions in this model also suggests a difference in the requisite structure of their biological counterparts. Certain implementations of global inhibition can act similarly to lateral inhibition in some ways (especially when the spatial propagation of global inhibition over time is modelled; see work by Sağlam and Hayashida [66]). However, the question of whether a single circuit element can be made to match the disparate requirements of the separate lateral and inhibitory circuits modelled here, and whether such a circuit is biologically supported, remain questions for future work.

When discussing the roles of global and lateral inhibition in dealing with wide-field motion, it should be noted that the model does not suppress such motion after looming has occurred, when only a small portion of the grating is visible near the edges of the eye. At this point not enough of the grating is exposed to form a coherent pattern, instead forming small repeated contrast transitions or translating elements, which generate a prominent LGMD response in vivo (e.g. O’Shea and Rowell [30] and Rowell et al. [14]). Biological data for stimuli matching this exact profile—that of a loom whose final visual subtense renders only a sliver of the visual field on the circumference of the eye visible—are not available from the literature. However, looming stimuli from Silva et al. [27], which were presented head-on to the animal, showed an extended decay phase in DCMD (and thus LGMD) firing after collision occurred when a grating background was present. It is possible that obstruction of a large portion of the visual field by the stimulus at collision interfered with wide-field inhibitory mechanisms in a similar way, and that this extended decay phase in firing is a related phenomenon to the post-collision firing seen here. This idea is further strengthened through the observation by Rowell et al. [14] that the LGMD produces increasing responses grating motion as the size of the grating region is reduced below 40°, implying that inhibitory mechanisms do not fully suppress such stimuli. Verifying that the post-collision activity of the model and the extended firing rate decay observed by Silva et al. [27] are related phenomena would require further testing of the model under additional stimuli and conditions.

While global and lateral inhibition contribute to control of dynamic range and suppression of wide-field motion responses in the model, the roles of direct feedforward inhibition to the LGMD are focused around peak time. Firstly, feedforward inhibition advances peak time relative to collision, an observation which agrees well with biological data [67]. This is important, as it is the timing of this peak—and the subsequent decay in firing rate—which controls the timing of escape jumps in locusts [5]. Given that removal of feedforward inhibition from the model reduced the time between LGMD peak firing and collision by approximately half for cases both with and without background motion, it is likely that feedforward inhibition is vital for ensuring that escape occurs early enough to be effective.

Aside from influencing when peak LGMD firing occurs during looms, feedforward inhibition also strongly affects its timing variability, with peak time becoming more consistent after its removal. While this is difficult to reference directly against experimental data, removal of feedforward inhibition by Gabbiani et al. [67] did substantially decrease variability in the timing of the last spike fired during looming responses, suggesting that the model behaviour may have biological relevance. As noted previously (see Simple looming responses), higher variability in peak times seems to correlate with increasing “burstiness” in the spike rasters. The mechanism for this appears to be that clustering spiking into prominent local maxima distributed around the average peak time increases the likelihood that the global maximum for a given run will be at one of these local maxima, offset from the average peak time. Given that Fig14A and B show similar differences in clustering on their rasters based on whether feedforward inhibition was present, there appears to be a strong case that feedforward inhibition controls peak time variability via clustering of spikes into bursts. With the OFF FFI neurons being relatively few in number—61 in total—in accordance with biological data [15], their resultant sparse spiking would tend to create specific windows for LGMD firing. This effect would be further reinforced by the included M currents, which are deactivated during inhibition-induced hyperpolarization, inducing a subsequent period of elevated resting potential and lengthened membrane time constant that creates enhanced integration of excitatory inputs and resultant rebound spiking. It would be valuable to determine whether the shaping of bursts by feedforward inhibition still occurs in the absence of M currents in the model.

Note that this mechanism of feedforward inhibitory input shaping LGMD bursts and peak time variation should mean that less clustering of spikes and variability in peak time should occur if inhibition were relayed more smoothly over time by a greater number of neurons. This agrees well with the reduced peak time variability for ON stimuli, as the greater number of ON FFI neurons that OFF (469 versus 61) and their weaker per-neuron input strength at the LGMD means temporally smoother inhibitory input over the course of an ON loom than OFF—an aspect of the model implemented based on biological observations [14]. Supplemental data from Dewell et al. [16] comparing OFF and ON loom responses from the LGMD do appear to show more prominent local maxima in the OFF responses, suggesting that the differences seen between these cases in the model may have a biological basis. Overall, a deeper comparison of both peak time variability and bursting properties between ON and OFF looms in vivo is justified, and would provide a valuable reference point for the model. Of particular use would be recent data from Mitra et al. [59], who compared key quantities of looming responses such as the timing and magnitude of peak firing rate between ON and OFF looms across multiple *l/* |*v*|, providing useful data for model fitting and comparison.

Achieving accuracy in the model’s relative responses to ON and OFF stimuli may require modelling the effects of light and dark adaptation which can occur prior to stimulus presentation, as this adaptation has effects on the temporal dynamics of photoreceptor responses in vivo [19]. Incorporating such elements would also allow investigation of the relative importance of these differences via the model itself.

While differences in between OFF and ON feedforward inhibition may explain the differences in peak time variability and apparent bursting properties between the corresponding looms, recall that the effects of field C excitation were also proposed as an explanation for reduced peak time variability in ON looms. These two explanations are not contradictory; however, further assessment of the model is needed to determine their relative contributions to this phenomenon.

While the LGMD bursting apparently created by feedforward inhibition seems to be the cause of its influence on peak time, the fact that the positioning of the OFF FFI neurons was varied each model run likely mediates the resultant increase in peak time variability. This has some significance regarding predictions that can be made for the LGMD in vivo. If the inhibitory neurons of the DUB shape bursting and peak time variability in the biological LGMD, then it is likely that the peak time varies

- from animal to animal; and
- with variations in input trajectory,

as these factors would vary the alignment of the looming stimulus with arrangement of underlying inhibitory DUB neurons. This provides another set of testable model predictions to examine in a study of bursting and peak time.

Based on the above, it could be inferred that the lack of peak time variability in ON stimuli is a result of not varying the positioning of the ON FFI neurons between model runs. This is likely partially true; however, the number of unique locations within the pattern of symmetry of the ON FFI neurons is only 5, as opposed to 19 for the OFF FFI neurons (see Fig 7 from Olson et al. [31]). This is a result of the greater density of neurons, and limits the possible variation in ON FFI neuron spike timing that different offsets can have.

When combined with the greater number of individually weaker inhibitory inputs limiting the possibility of clustering LGMD spiking between them, it seems unlikely that ON FFI neurons are capable of generating similar peak time variability—though further investigation of the model in this regard is still warranted.

While the model suggests that feedforward inhibition contributes variability to the peak time of the LGMD looming response, this raises the question of what the importance of this variability is. In locusts, the greater variability in peak time seen at higher *l/* |*v*| looms [1] corresponds directly to greater variability in escape behaviour timing seen at higher *l/* |*v*| values [68]. Thus, a distinct possibility is that feedforward inhibition-induced peak time variability serves to make escape behaviour less predictable, increasing the chance of avoiding predation. Based on factors discussed above, this variability would emerge most strongly between individuals and between different approach trajectories; this would align with other aspects of locust behaviour which seem to reduce predictability across differing individuals and approach directions, such as lateralization [69].

The roles of the different inhibitory mechanisms in the model can be summarized as follows:

- lateral inhibition inhibits responses to wide-field motion, particularly sustained ones;
- global inhibition controls dynamic range by curtailing excitation near to collision, reduces initial responses to wide-field motion, and improves distinguishing of looming stimuli against moving backgrounds; and
- feedforward inhibition shapes LGMD peak firing rate during looming, resulting in an earlier—and in the case of OFF feedforward inhibition, more variable—peak time.

The above functions relate to stimuli which the model reproduces well on various qualitative and quantitative levels; as such, this strengthens the case that such mechanisms play similar roles in the LGMD circuit in vivo.

### Issues with coherence selectivity

The most visible way in which the model differs from the biological LGMD is in its lack of coherence selectivity, to the extent of having a preference for *incoherent* stimuli. The clearest potential cause for this is the omission of a key mechanism of coherence selectivity in vivo: lateral excitation. Blocking of lateral excitation in vivo via scopolamine removes the majority of coherence dependence in LGMD OFF loom responses, though it does not do so completely [33]. Thus, while the omission of lateral excitation explains why the model is not selective for coherence, it does not explain why it prefers incoherent stimuli.

Observations of the model with only the direct inputs to the field A of LGMD rendered spatially incoherent were included in this work to determine what level of coherence selectivity is implemented at the LGMD itself. Biologically speaking, coherence selectivity at this level of the system is understood to arise from active conductances in field A—namely, the HCN and K_D-like_ conductances. However, despite inclusion of these conductances, the modelled LGMD shows no distinct coherence preference, aside from a slightly greater response to incoherent OFF looms around the firing rate peak. As mentioned previously (see Stimulus coherence), a purely passive LGMD should prefer incoherent stimuli throughout looming [32], so the fact that field A of the model has a largely neutral coherence preference suggests that active conductances may be partially effective as-implemented.

However, both the experimental data and modelling work of Dewell and Gabbiani [32] suggest that these active conductances should confer greater coherence selectivity. The reason they do not in this model may be due to various factors. One of these is the mathematics used for the HCN conductance. While the parameters and equations used were based on intracellular LGMD recordings and subsequent analysis by Dewell et al. [32], the limitations imposed by their recording methods mean that the actual channel kinetics may differ somewhat. It is notable that the LGMD model presented by Dewell et al. in the same work used a slightly different voltage dependency of HCN channel time constant than that derived directly from their experimental data, and was able to produce coherence selectivity comparable to that seen biologically. As such, varying the dynamics of the HCN channels in the model presented here is a promising avenue for improving the current model’s biological fidelity.

Another possible reason that the active conductances do not produce greater coherence selectivity in the present model relates not to the conductances themselves, but to the geometry and input mapping of field A. The coherence selectivity conferred by HCN and K_D-like_ conductances is contingent on generating sufficient localized increases in membrane voltage to create channel activations and inactivations. However, as seen for the reduced example in Fig 2, the distalmost branch level accounts for approximately half of the eye. As such, each distal branch segment spans nearly half the eye in terms of ommatidia, but is very narrow. The narrowness in terms of retinotopic units means limited inputs arrive at each distal branch segment, while the representation of each of these segments as a single compartment in the model means that inputs are effectively spread out over the entire segment length. This limits the localization of input needed for the active conductances to function. Conversely, inputs arriving in more proximal branch segments create different difficulties. These proximal segments have increased intercompartmental conductances and increased capacitances due to their higher diameters—both of which limit local voltage increases during excitatory input. These problems resulting from model geometry would become more prominent later during looming, as the edge of the stimulus spreads to ommatidia that map to more proximal and distal sites in field A. This corresponds well to the phenomenon observed in Figs. 8, 11 and 10, wherein the lowest coherence selectivity (or greatest incoherence selectivity) for OFF looms occurs late in the loom, near the LGMD firing peak; such correspondence lends this explanation credence.

Despite the omission of lateral excitation and the limited efficacy of active field A conductances, the model does exhibit coherence preference when lateral inhibition is removed. This demonstrates two things:

- that lateral inhibition confers a preference for incoherent stimuli; and
- that some other mechanism of creating coherence selectivity is present in the model.

The first point is unsurprising, given that lateral excitation is one of the primary methods of generating coherence preference in the LGMD [33]. However, given that lateral inhibition is a motif within the LGMD circuit [30], it places increased importance on the existing mechanism of coherence selectivity within the LGMD and its presynaptic network, as these must not only generate coherence preference, but also first overcome a preference for incoherence. The second point is somewhat more surprising, as only lateral excitation and active field A conductances have been described previously as mechanisms of promoting coherence selectivity in vivo.

Based on examinations of FFI neuron responses to varying stimulus coherence, it seems extremely likely that feedforward inhibition is the primary mechanism creating coherence selectivity in this model. The modelled FFI neurons exhibit a preference for incoherent stimuli, even when such preferences are removed from the inputs they share with the LGMD. This represents a completely different means of implementing coherence selectivity in the LGMD: instead of providing boosted excitation with increasing coherence, the modelled FFI neurons provide inhibition that increases as coherence decreases. An interesting facet of this is that the coherence preference provided by the FFI neurons to the LGMD increases as the coherence preference of their shared inputs decreases. This suggests that it has value as a compensatory mechanism for the incoherence preference created by lateral inhibition, which is otherwise a useful element in the model.

To allow the model to reproduce biological levels of coherence selectivity, multiple avenues are available based on the observations above. These include

- implementing lateral excitation between TmA inputs, based on data from Zhu et al. [33];
- modifying the compartmentalization of field A to incorporate greater subdivision of distal branch segments;
- modifying the input mapping of field A to exclude the most proximal compartments;
- modifying the dimensions of field A to reduce the diameter of more proximal compartments, decreasing their capacitance and axial conductance; and
- re-examining the presynaptic network of the LGMD to determine whether lateral inhibition can be further reduced while still performing its function.

Examining these various avenues may yield further insight on the function and relative importance of the different mechanisms of generating coherence selectivity in the biological LGMD.

### Translating and transitioning stimuli

While the model captures the looming preference of the LGMD over the trajectories tested, along with the capacity of the LGMD to rapidly transition between response trajectories when stimulus behaviour changes [1, 23], certain subtle differences between the model and its biological counterpart gives clues as to where improvement is possible. The discrepancies in experimental and model firing rates during translation can be explained in the context of the model structure (see Fig 2) and input mapping (see Fig 4). As discussed above, almost half of the eye on the distalmost side was mapped to the distalmost branching level of field A, with each branch segment therein represented by a single compartment. This means that inputs ranging from the anteriormost edge of the visual field to near the center are all mapped to a compartment at the same electrotonic distance from the LGMD SIZ. This might explain the model’s initial overestimation of firing rate when the translating stimulus is nearer to the anterior edge of the eye, followed by underestimation as it draws closer to the center of the visual field; in vivo, such inputs would become stronger as they draw closer to the SIZ and are subject to less attenuation. Later during the simulated translation, the leading edge of the stimulus eventually crosses into regions mapped to a more proximal branching level of field A, where the instantaneous increase in diameter and decrease in distance from the SIZ causes a large drop in electrotonic distance between synaptic inputs and the SIZ. This would occur shortly before the stimulus crosses the central axis of the modelled eye, and explains the sudden increase in firing rate seen around this time. The overestimation of firing rate later in the course of translation suggests that model inputs to the proximal portions of field A are at an insufficient electronic distance from the SIZ. This could either mean that inputs should not be mapped to the most proximal level (or levels) of field A, or that these levels account for an insufficient portion of the electrotonic size of field A; more work is needed to determine this.

Note that the structural issues discussed above as possible causes for discrepancies in the translating response—namely, insufficient subdivision of distal branch levels, and mapping input to proximal levels with insufficient electrotonic separation from the SIZ—are identical to those put forth as reasons why active conductances failed to generate coherence selectivity in field A (see Issues with coherence selectivity). Given that these factors are a possible explanation for multiple discrepancies between model and biological behaviour, addressing them should be a high priority going forward.

The inconsistencies between model translation responses and biological data may also partially explain the lack of a characteristic drop in firing rate at the transition to looming. The underestimation of LGMD firing rate over most of the translating phase would reduce the buildup of inhibitory *g*_*A*_ currents that may contribute to the drop in firing rate after transition. Furthermore, the sudden increase in excitatory drive as the stimulus approaches the centerline of the eye, and excitation is mapped to more proximal sites in field A, may have masked an expected drop in firing rate that would otherwise occur. This requires further investigation, possibly by shifting the transition from translation to looming further posterior within the visual field, away from the transition out of the distalmost portion of field A.

It should be noted that, to our knowledge, this represents the first time in the literature an LGMD model has been subjected to such a range of stimulus trajectories. While both this model and prior models elsewhere in the literature [18, 32] have faithfully reproduced looming responses, and both this model and prior models [36, 70] have compared looming and translating responses, this modelling work has been the first to examine transitions between looming and translation, with their accompanying signature effects [23] on LGMD spiking. As such, it is unclear whether the aspects of the model’s responses to complex stimuli which do not resemble biological data are a consequence of structural elements peculiar to this model alone, or whether other LGMD models exhibit similar behaviour.

### Future directions

The most direct next step from the work presented here is to find and address the causes of observed discrepancies between model behaviour and biological data. As noted above for some cases, adjusting single elements of the model may be able to address multiple observed issues.

Further insights can also be gained by examining aspects of the model not explored here. One particularly notable aspect is bursting; while qualitative examinations of spike raster data have enabled a cursory discussion of bursting in this work, performing a quantitative analysis of bursting would enable deeper comparisons with available literature data (e.g. McMillan and Gray [23], and Dewell and Gabbiani [38]). As bursting behaviour in the model appeared to vary in interesting ways as a function of stimulus and model structure, confirming that baseline bursting behaviour during looming is biologically representative would give observations from the model greater weight.

Another aspect of the model to consider is its response to stimuli which instantaneously change their approach speed instead of their direction of motion. While in vivo responses to these stimuli [1] have elements in common with transitions to and from looming trajectories [23], certain aspects differ. Comparing responses to this stimulus type to responses to trajectory changes examined here could illuminate subtle ways in which the model differs from biology—especially if the issues currently present in translation and translation-to-looming responses can be addressed first.

Looking beyond adjustments to the model in its current scope, future work should look to incorporate modelling of the locust visual system into models of *visuomotor* systems. This presents the ultimate test of any visual model: does it produce responses which encode signals for biologically-appropriate motor behaviour? Some reference data exists for such investigations, such as

- how different phases of the LGMD response should shape preparation for and triggering of escape jumps [5];
- how changing stimulus parameters (such as coherence and ON vs OFF) change jump probabilities [16]; and
- how DCMD firing relates to evasive gliding in flight [71].

Potential future work of this type also provides the impetus for a model which seeks to capture responses to a diverse array of stimuli, as it improves comparability with potential closed-loop examinations of locust behaviour. Animals permitted to interact with stimuli in a closed-loop fashion can modify these stimuli in unexpected ways, and thus attempting to replicate the responses requires models that can be expected to respond realistically to a variety of stimuli. Overall, while the model presented here attempts to capture a wide scope in terms of both elements included and responses replicated, further work is required.

## Notes

### Competing Interest Statement

The authors have declared no competing interest.

